# Structure-informed microbial population genetics elucidate selective pressures that shape protein evolution

**DOI:** 10.1101/2022.03.02.482602

**Authors:** Evan Kiefl, Ozcan C. Esen, Samuel E. Miller, Kourtney L. Kroll, Amy D. Willis, Michael S. Rappé, Tao Pan, A. Murat Eren

## Abstract

Comprehensive sampling of natural genetic diversity with metagenomics enables highly resolved insights into the interplay between ecology and evolution. However, intra-population genomic variation represents the outcome of both stochastic and selective forces, making it difficult to identify whether variants are maintained by adaptive, neutral, or purifying processes. This is partly due to the reliance on gene sequences to interpret variants, which disregards the physical properties of three-dimensional gene products that define the functional landscape on which selection acts. Here we describe an approach to analyze genetic variation in the context of predicted protein structures, and apply it to study a marine microbial population within the SAR11 subclade 1a.3.V, which dominates low-latitude surface oceans. Our analyses reveal a tight association between the patterns of nonsynonymous polymorphism, selective pressures, and structural properties of proteins such as per-site relative solvent accessibility and distance to ligands, which explain up to 59% of genetic variance in some genes. In glutamine synthetase, a central gene in nitrogen metabolism, we observe decreased occurrence of nonsynonymous variants from ligand binding sites as a function of nitrate concentrations in the environment, revealing genetic targets of distinct evolutionary pressures maintained by nutrient availability. Our data also reveals that rare codons are purified from ligand binding sites when genes are under high selection, demonstrating the utility of structure-aware analyses to study the variants that likely impact translational processes. Overall, our work yields insights into the governing principles of evolution that shape the genetic diversity landscape within a globally abundant population, and makes available a software framework for structure-aware investigations of microbial population genetics.

**Significance:** Increasing availability of metagenomes offers new opportunities to study evolution, but the equal treatment of all variants limits insights into drivers of sequence diversity. By capitalizing on recent advances in protein structure prediction capabilities, our study examines subtle evolutionary dynamics of a microbial population that dominates surface oceans through the lens of structural biology. We demonstrate the utility of structure-informed metrics to understand the distribution of nonsynonymous polymorphism, establish insights into the impact of changing nutrient availability on protein evolution, and show that even synonymous variants are scrutinized strictly to maximize translational efficiency when selection is high. Overall, our work illustrates new opportunities for discovery at the intersection between metagenomics and structural bioinformatics, and offers an interactive and scalable software platform to visualize and analyze genetic variants in the context of predicted protein structures and ligand-binding sites.

## Introduction

Genetic diversity within populations emerges from and is shaped by a combination of stochastic and selective pressures, which often lead to phenotypic differences between closely related individuals, sometimes within a few generations (Burke et al. 2010; Lenski et al. 1991). Surveys of microbial communities within natural habitats through phylogenetic marker genes (Olsen et al. 1986; Acinas et al. 2004; Sogin et al. 2006) and metagenomics (Simmons and DiBartolo et al. 2008; Allen et al. 2007) have revealed a tremendous amount of genetic variation within environmental populations (T. P. Curtis and Sloan 2005; Thomas P. Curtis et al. 2006), and an ever-increasing number of available genomes and metagenomes have provided insight into the selective pressures that shape such variation. However, the overwhelming complexity and dynamicity of these selective pressures, which occur even in the simplest environments (Good et al. 2017), has hindered our ability to determine which variants are under the influence of which pressures (Ochman 2003; Mes 2008).

Inferring selective pressures through the isolation of microbial strains and comparative genomics has been widely successful. More recently, metagenome-assembled genomes (L.-X. Chen et al. 2020) and single-amplified genomes (Woyke, Doud, and Schulz 2017) have dramatically increased the number (Almeida et al. 2021; Pachiadaki et al. 2019; Paoli et al. 2021) and diversity (Hug et al. 2016) of microbial clades represented in genomic databases, offering an even larger sampling of environmental microbes to study the emergence and maintenance of genetic variation (Garud and Pollard 2020). Nevertheless, genomes represent static snapshots of individual members of often complex environmental populations, and thus, working with genomic sequences alone substantially undersamples genetic variability in natural habitats and its associations with environmental and ecological forces (Van Rossum et al. 2020). This shortcoming is partially addressed by shotgun metagenomics (Quince et al. 2017) and metagenomic read recruitment, where environmental sequences that are aligned to a reference can be studied to identify genetic variants at the resolution of single nucleotides (Whitaker and Banfield 2006; Denef 2019). In particular, using genomes to recruit reads from metagenomes enables a comprehensive sampling of all genetic variants within environmental populations (Simmons and DiBartolo et al. 2008). Due to the immensity of sequencing data generated by metagenomic studies, even subtle genetic variation in natural populations is now resolvable, making it possible to explicitly correlate patterns of genomic variation with temporal or spatial environmental variables to elucidate the interplay between ecology and evolution (Schloissnig et al. 2013; Bendall et al. 2016; Anderson et al. 2017; Delmont et al. 2019; Garud et al. 2019; Zhao et al. 2019; Shenhav and Zeevi 2020; Olm et al. 2021; Conwill et al. 2022). Although quantification and analysis of sequence variants derived from metagenomic data has improved dramatically, inferring the functional impact of individual nucleotides remains a fundamental challenge in part due to the sole reliance on DNA sequences, which do not represent physical properties of proteins they encode, and thus disguise the functional impact of individual mutations.

Given the intermediary role that structure plays within the ‘sequence-structure-function paradigm’ (Anfinsen 1973), including protein structures as a dimension of analysis is commonplace in studies of protein evolution (Siltberg-Liberles, Grahnen, and Liberles 2011; Harms and Thornton 2013; Sikosek and Chan 2014), and it is appreciated that the accuracy of evolutionary models improves with combined analyses of protein structures and the evolution of underlying sequences (Wilke 2012). In contrast, the state-of-the-art approaches that quantify genetic variants in environmental microbial populations typically treat genes as strings of nucleotides (Schloissnig et al. 2013; Eren et al. 2015; Nayfach et al. 2016; Costea et al. 2017; Olm et al. 2021). While this strategy enables rapid surveys of population dynamics through single-nucleotide variants, it disregards the physical properties of three-dimensional gene products that selection acts upon, and thus misses a critical intermediate to understand the relationship between selection and fitness (Golding and Dean 1998; K. Chen and Arnold 1993). The importance of mapping sequence variants on predicted protein structures to identify genetic determinants of phenotypic variation has been noted more than two decades ago (Sunyaev, Lathe, and Bork 2001), yet the limited availability of protein structures has historically rendered protein structure-informed microbial population genetics impractical. Given dramatic advances in both solving and predicting protein structures in recent years (Kuhlman and Bradley 2019), most notably deep learning approaches such as AlphaFold (Jumper et al. 2021) that offer highly accurate protein structure predictions, this constraint is likely a problem of the past. Altogether, open questions in microbial ecology and evolution, advances in computation, and increased availability of data are culminating in a research landscape that is ripe for new software solutions that integrate protein structures with ’omics data in order to observe and interpret evolutionary processes that shape sequence variation in natural populations.

Here we develop an interactive and scalable software solution for the analysis and interactive visualization of metagenomic sequence variants in the context of predicted protein structures and ligand binding sites as a new module in anvi’o, an open-source, community-led multi-omics platform (https://anvio.org). By importing AlphaFold-predicted protein structures into *anvi’o structure*, we (1) demonstrate the shortcomings of purely sequence-based approaches to interpret patterns of polymorphism observed within complex microbial populations, (2) propose two structural features to interpret genetic variation, RSA and DTL, (3) illustrate that nonsynonymous polymorphism is more likely to encroach upon active sites when selection is low, but is purged from active sites when selection is high, and (4) provide evidence that common codons are more translationally robust than their rare synonymous counterparts, which appear within structurally/functionally noncritical sites when selection is low.

## Results and Discussion

To investigate selective pressures that drive protein evolution within microorganisms inhabiting complex naturally occurring environments, we chose a single microbial taxon and a set of metagenomes that match to its niche boundaries: SAR11 (*Candidatus* Pelagibacter ubique), a microbial clade of free-living heterotrophic alphaproteobacteria that dominates surface ocean waters (Morris et al. 2002), and Tara Oceans Project metagenomes (Sunagawa et al. 2015), a massive collection of deeply sequenced marine samples from oceans and seas across the globe. SAR11 is divided into multiple subclades with distinct ecology (Giovannoni 2017). Thus, we further narrowed our focus to HIMB83, a single SAR11 strain genome that is 1.4 Mbp in length. HIMB83 is a member of the environmental SAR11 lineage 1a.3.V, one of the most abundant (Nayfach et al. 2016) and most diverse (Delmont & Kiefl et al. 2019) microbial lineages in marine systems, which recruits as much as 1.5% of all metagenomic short reads in surface ocean metagenomes (Delmont & Kiefl et al. 2019).

To quantify the genetic variability of 1a.3.V, we used HIMB83 as a reference genome of the subclade, and competitively recruited short reads (see Methods) from 93 low-latitude surface ocean metagenomes (Table S1), resulting in 390 million reads that were 94.5% identical to HIMB83 on average (Figure S1). As an individual member of a diverse subclade, HIMB83 possesses a genomic context that is insufficient for resolving the extent of genetic diversity within 1a.3.V. Regardless, HIMB83 possesses the ’core’ gene set of 1a.3.V, and so reads recruited by these genes represent the diversity of the 1a.3.V core genome. Of the 1,470 genes in HIMB83, we restricted our analysis to 799 genes that we determined to form the 1a.3.V core genes, and 74 metagenomes in which the average coverage of HIMB83 exceeded 50X (see Methods). The reads recruited to the 1a.3.V core represent a dense sampling of the diversity within this environmental lineage that far exceeds the evolutionary resolution and volume of sequence data achievable through comparisons of cultured SAR11 genomes alone (Figure S1). As a result, these data provide a unique opportunity to zoom in and track how genomic variation in one of the most abundant microbial populations on Earth shifts in response to ecological parameters throughout the global ocean (Figure S2).

### Polymorphism rates reveal intense purification of nonsynonymous mutants

To quantify genomic variation in 1a.3.V, in each sample we identified codon positions of HIMB83 where aligned metagenomic reads did not match the reference codon. We considered each such position to be a *single codon variant* (SCV). Analogous to single nucleotide variants (SNVs), which quantify the frequency that each nucleotide allele (A, C, G, T) is observed in the reads aligning to a nucleotide position, SCVs quantify the frequency that each codon allele (AAA, …, TTT) is observed in the reads aligning to a codon position (see Methods for a more complete description). Since SCVs are defined to be ‘in-frame’, they provide inherent convenience when relating nucleotide variation in the genomic coordinates to amino acid variation in the corresponding protein coordinates, as well as for determining whether or not nucleotide variation leads to synonymous or nonsynonymous change. Within the 1a.3.V core genes, we found a total of 9,537,022 SCVs, or 128,879 per metagenome on average. These SCVs distributed throughout the genome such that 78% of codons (32% of nucleotides) exhibited minor allele frequencies >10% in at least one metagenome. Despite this extraordinary level of diversity, our read recruitment strategy is stringent and yields reads that on average differ from HIMB83 in only 6 nucleotides out of 100 (Table S2), precluding the possibility that this diversity is generated from excessive nonspecific mapping. While puzzling, this level of diversity is not surprising as it agrees with numerous studies that have pointed out the astonishing complexity of the SAR11 subclade 1a.3.V (Nayfach et al. 2016; Delmont and Kiefl et al. 2019; Haro-Moreno et al. 2020) that could not be further divided into sequence-discrete populations (Delmont and Kiefl et al. 2019).

We found this diversity to be overwhelmingly synonymous. By splitting each SCV into its synonymous (s) and nonsynonymous (ns) proportions, we calculated per-site rates of s- polymorphism and ns-polymorphism as pS^(site)^ and pN^(site)^, not to be confused with the related concepts dS and dN. While dS and dN quantify rates of synonymous and nonsynonymous *substitution* between diverged species, pN^(site)^ and pS^(site)^ can (1) resolve shorter evolutionary timescales than the characteristic fixation rate, (2) be calculated from metagenomic read recruitment data without complete haplotypes, and (3) define rates on a per-sample basis, thus enabling inter-sample comparisons. Overall, we found that the average pS^(site)^ outweighed pN^(site)^ by 19:1 (Table S3), revealing an overwhelming fraction of the 1a.3.V diversity to be synonymous and illustrating how nonsynonymous mutants are purified at a much higher rate than synonymous mutants in the population at large.

### Nonsynonymous polymorphism avoids buried sites

pN^(site)^ values varied significantly from site-to-site and from sample-to-sample, but overall, more variation existed between sites in a given sample than between samples of a given site (Figure S3). The extent that a given site can tolerate ns-polymorphism is largely determined by the local physicochemical environment of the encoded residue, which is defined by the 3D structure of the protein. Thus, we broadened our focus by developing a computational framework, *anvi’o structure* (Supplementary Information), that enabled the integration of environmental sequence variability with predicted protein structures (Figure 1).

**Figure 1.**
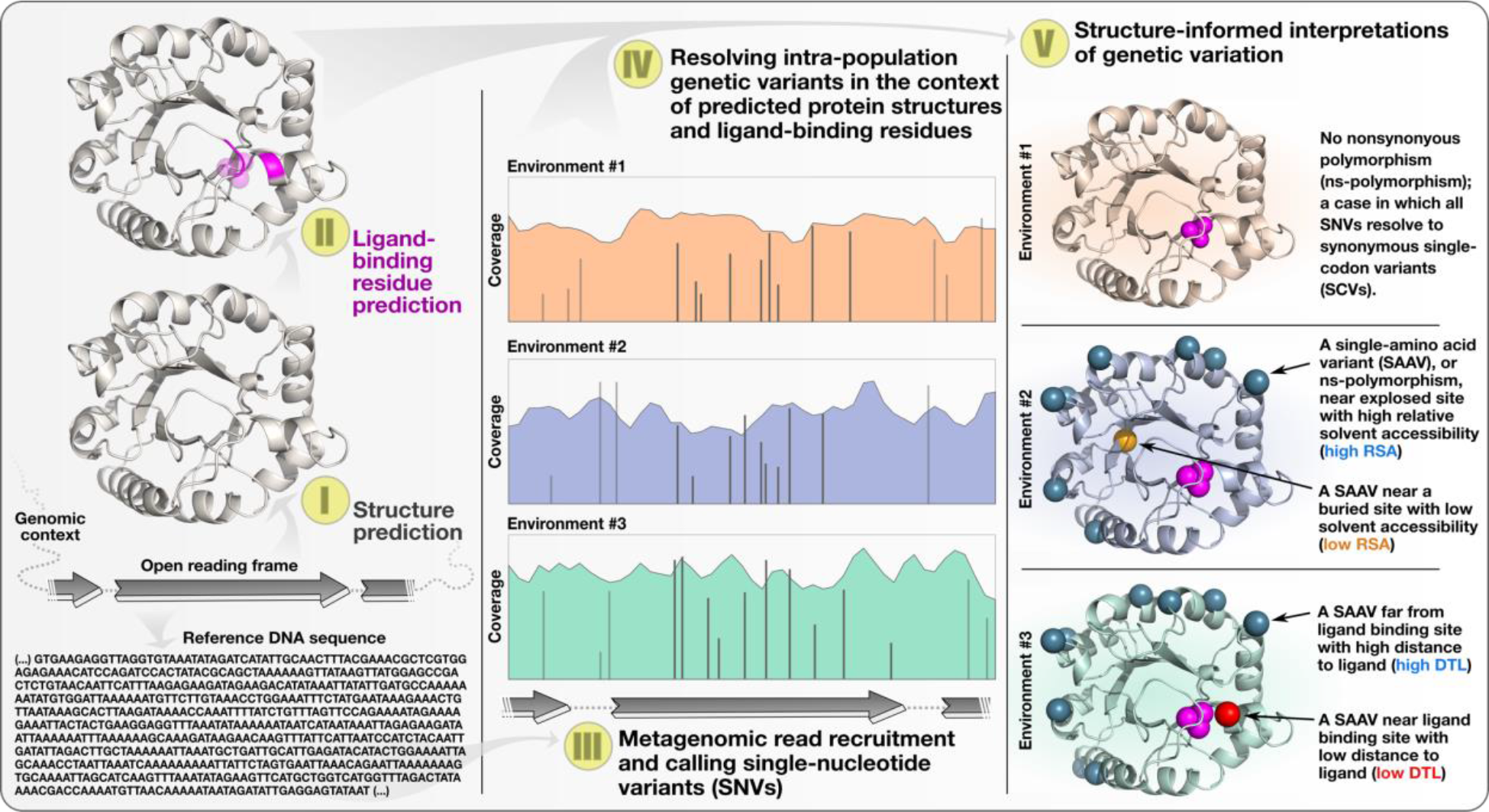
Anvi’o workflow for structure-informed population genetics.

We used two independent methods to predict protein structures for the 799 core genes of 1a.3.V: (1) a template-based homology modeling approach with MODELLER (Webb and Sali 2016), which predicted 346 structures, and (2) a transformer-like deep learning approach with AlphaFold (Jumper et al. 2021), which predicted 754. Our evaluation of the 339 genes for which both methods predicted structures (Supplementary Information) revealed a comparable accuracy between AlphaFold and MODELLER (Figure S4, Table S4). Thus, we opted to use AlphaFold structures for all downstream analyses due to its higher structural coverage. Indeed, AlphaFold - predicted protein structures covered over 90% of the core genes, highlighting the emerging opportunities afforded by recent advances in de novo structure prediction.

Aligning single-codon variants to predicted structures enabled us to directly compare the distributions of s-polymorphism and ns-polymorphism rates relative to biophysical characteristics of the encoded proteins. We first investigated the association between polymorphism rates and relative solvent accessibility (RSA), a biophysical measure of how exposed (RSA = 1) or buried (RSA = 0) a site is. Since nonsynonymous mutations at buried sites are more likely to disrupt folding and stability, RSA serves as a powerful proxy to discuss the strength of structural constraints acting at a site (Echave, Spielman, and Wilke 2016). By calculating RSA for each site in the predicted structures, and then weighting every site by the pN^(site)^ and pS^(site)^ across all samples, we established proteome-wide distributions for pN^(site)^ and pS^(site)^ relative to RSA (Figure 2a). These data showed that pS^(site)^ closely resembled the null distribution, which illustrates the lack of influence of RSA on s-polymorphism, while pN^(site)^ deviated significantly and instead exhibited strong preference for sites with higher RSA. This finding aligns well with the expectation that buried sites are likely to purify nonsynonymous change due to disruption of protein stability while being relatively more tolerant to synonymous change, and validates our methodology.

**Figure 2.**
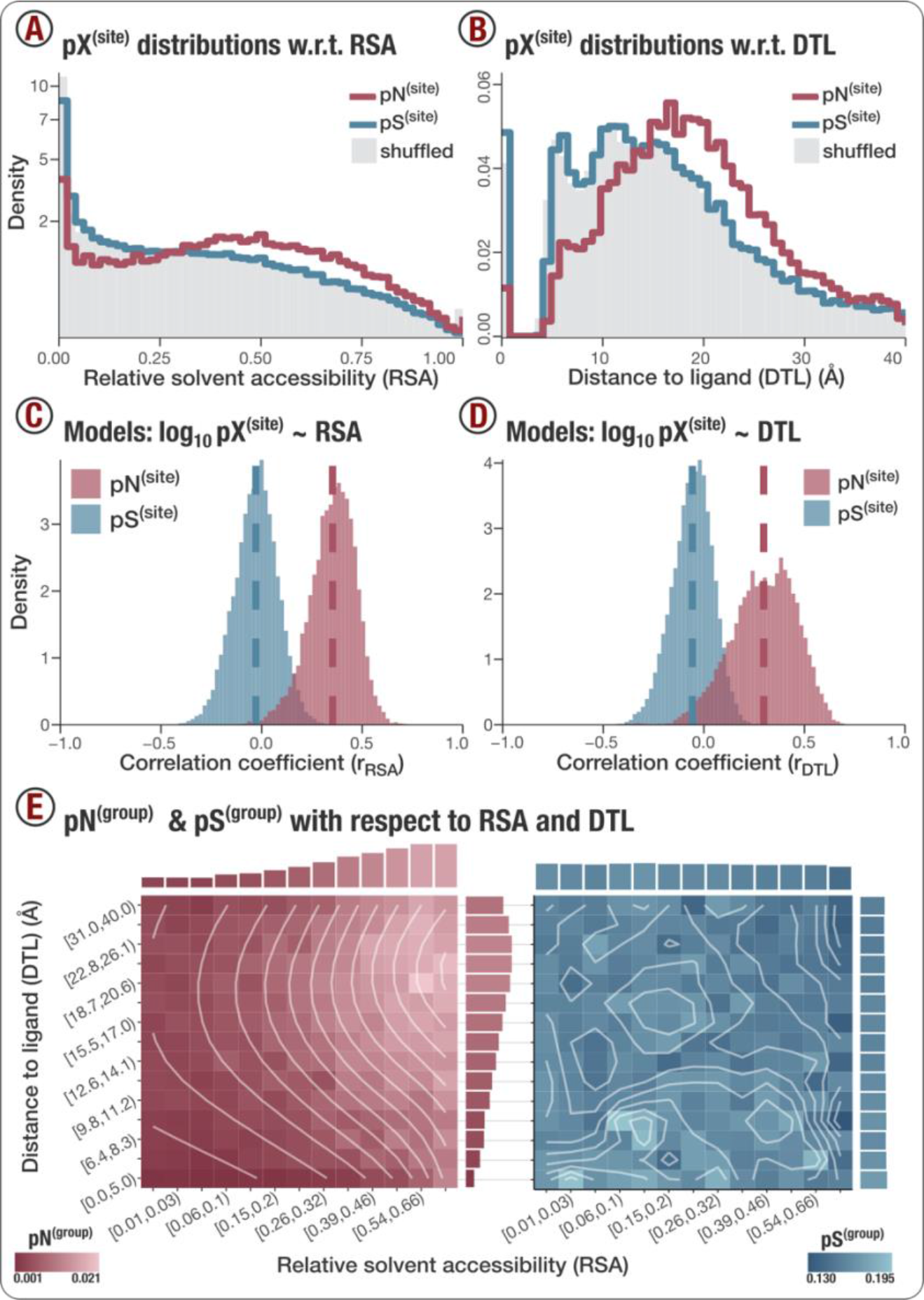
(A) Structural constraints shift the pN^(site)^ distribution towards high relative solvent accessibility (RSA). The pN^(site)^ distribution (red line) and pS^(site)^ distribution (blue line) were created by weighting the RSA values of 239,528 sites (coming from the 754 genes with predicted structures) by the pN^(site)^ and pS^(site)^ values observed in each of the 74 samples, totaling 17,725,072 pN^(site)^ and pS^(site)^ values. The average distribution of 10 independent, randomly shuffled datasets of pN^(site)^ is depicted by the grey- regions for pN^(site)^, and represents the null distribution expected if no association between pN^(site)^ and RSA existed. Since the null distribution for pS^(site)^ so closely resembles the null distribution for pN^(site)^, it has been excluded for visual clarity, but can be seen in Figure S5. **(B) Functional constraint shifts the pN^(site)^ distribution towards high distance-to-ligand (DTL) values.** The pN^(site)^ distribution (red line) and pS^(site)^ distribution (blue line) were created by weighting the DTL values of 155,478 sites (coming from 415 genes that had predicted structures and at least one predicted ligand) by the pN^(site)^ and pS^(site)^ values observed in each of the 74 samples, totaling 11,505,372 pN^(site)^ and pS^(site)^ values. The pN^(site)^ null distribution was calculated according to the procedure described in panel A, where again, the pS^(site)^ null distribution closely resembled the pN^(site)^ null distribution, and can be seen in Figure S5. **(C) Linear models reveal positive correlations between pN^(site)^ and RSA.** The two distributions show Pearson correlation coefficients produced by linear models of the form log10(pN^(site)^) ∼ RSA (red-filled region) and log10(pS^(site)^) ∼ RSA (blue-filled region). A model has been fit to each gene-sample pair that passed filtering criteria (see Supplementary Information), resulting in 16,285 nonsynonymous models and 24,553 synonymous models. Distribution means are visualized as dashed lines. **(D) Per-group polymorphism rates explain the major selective pressure trends with respect to RSA and DTL.** The left and right panels show heatmaps of pN^(group)^ and pS^(group)^. Each cell represents a group defined by RSA and DTL ranges shown on the x- and y- axes, respectively. The color of each cell represents the respective value for the group, where dark refers to low values and light refers to high values. White lines show the contour lines of smoothed data.

### Nonsynonymous polymorphism avoids active sites

While structural constraints ensure a given protein folds properly and remains stable, they do not guarantee its function. Comprehensive analyses of diverse protein families show that residues that bind or interact with ligands are depleted of mutations (Kobren and Singh 2019) due to strong selective pressures that maintain active site conservancy. This constraint is not limited to the immediate vicinity of ligand-binding residues, and has been observed to radiate outwards from the active site with a strength inversely correlated with distance from active site (Dean et al. 2002). We considered this distance as the ‘distance-to-ligand’ (DTL), and hypothesized that DTL may be a suitable proxy for investigating functional constraints in a manner complementary to RSA, a proxy for investigating structural constraints. To test this, we investigated distributions of pN^(site)^ and pS^(site)^ as a function of DTL for each predicted structure by first predicting sites implicated in ligand binding using InteracDome (Kobren and Singh 2019), and then calculating a DTL for each site, given the closest predicted ligand-binding site (Table S5).

The average per-site ns-polymorphism rate throughout the 1a.3.V core genome was 0.0088, however, we observed a nearly 4-fold reduction in this rate to just 0.0024 at predicted ligand binding sites (DTL = 0), indicating stronger purifying selection at ligand-binding sites (Figure 2b). Sites neighboring ligand-binding regions also harbored disproportionately low rates of ns- polymorphism, as indicated by the significant deviation towards larger DTL values. This illustrates that purifying selection that preserves proper ligand-binding functionality is not limited to residues at ligand-binding sites, but extends to proximal sites as well. When we defined DTL in sequence space rather than Euclidean space, this effect was no longer observable beyond sequence distances of ∼5-10 amino acids (Figure S6). Comparatively, pS^(site)^ deviated minimally from the null distribution. Overall, integrating predicted protein structures and ligand-binding sites into the analysis of the genetic diversity of an environmental population has enabled us to demonstrate that (1) structural constraints bias pN^(site)^ distributions towards solvent exposed sites (*i.e.* high RSA) (Figure 2a), and (2) functional constraints bias pN^(site)^ distributions towards sites that are distant from ligand-binding sites (*i.e.* high DTL) (Figure 2b).

### Proteomic trends in purifying selection are explained by RSA and DTL

Given the clear shift in ns-polymorphism rates towards high RSA and DTL sites across genes, we next investigated the extent that RSA and DTL can predict per-site polymorphism rates. By fitting a series of linear models to log-transformed polymorphism data (Table S6), we conclude that RSA and DTL can explain 11.83% and 6.89% of pN^(site)^ variation, respectively. Based on these models we estimate that for any given gene in any given sample, (1) a 1% increase in RSA corresponds to a 0.98% increase in pN^(site)^, and (2) a 1% increase in DTL (normalized by the maximum DTL in the gene) corresponds to a 0.90% increase in pN^(site)^. In a combined model, RSA and DTL jointly explained 14.12% of pN^(site)^ variation, and after adjusting for gene-to-gene and sample-to-sample variance, 17.07% of the remaining variation could be explained by RSA and DTL. In comparison, only 0.35% of pS^(site)^ variation was explained by RSA and DTL. Using a complementary approach, we constructed models for each gene-sample pair (Supplementary Information), the correlations of which we used to visualize the extent that pN^(site)^ can be modeled by RSA and DTL relative to pS^(site)^ (Figures 2c, 2d). Analyzing gene-sample pairs revealed that the extent of ns-polymorphism rate that can be explained by RSA and DTL is not uniform across all genes (Table S7) and can reach up to 52.6% and 51.4%, respectively (Figures S7, S8). Finally, we averaged polymorphism rates within groups of sites that shared similar RSA and DTL values, which demonstrated the tight association between the rate of within population ns-polymorphism rate and protein structure (Table S8, Figure 2e). Linear regressions of these data show that 83.6% of per-group ns- polymorphism rates and 20.7% of per-group s-polymorphism rates are explained by RSA and DTL (Supplementary Information).

The true predictive power of RSA and DTL for polymorphism rates is most likely higher than we report, since our approaches suffer from methodological shortcomings. For instance, we calculate RSA from the steric configurations of residues in predicted structures. Thus, errors in structure prediction propagate to errors in RSA. Errors in structure also propagate to errors in DTL, since DTL is calculated using Euclidean distances between residues, which is exacerbated by the uncertainty associated with ligand-binding site predictions. Furthermore, RSA and DTL calculations assume that the protein is monomeric, even though oligomeric proteins are common, and they represent the majority of proteins in some organisms (Goodsell and and Olson 2003). In these cases, exposed sites in the monomeric structure could be buried once assembled into the quaternary structure, and this is similarly true for estimates of DTL. Even if we assume structural predictions are 100% accurate, it is notable that binding site predictions exclude (1) ligands that are proteins, (2) ligand-protein complexes that have not co-crystallized with each other, (3) ligands of proteins with no shared homology in the InteracDome database, and (4) unknown ligand-protein complexes. Each of these shortcomings leads to missed binding sites, which leads to erroneously high DTL values in the proximity of unidentified binding sites (Figure S9). Furthermore, our predictions assume that if a homologous protein in the InteracDome database binds to a ligand with a particular residue, then so too does the corresponding residue in the HIMB83 protein. This leads to uncertain predictions, since homology does not necessitate binding site conservancy. Yet, despite these methodological shortcomings, our analyses show that RSA and DTL are significant predictors of per-site and per-group variation.

Clear partitioning of environmental genetic variation by RSA and DTL (Figure 2) highlights the utility of these metrics for studies of evolution following the increasing availability of protein structures. Analyses of total genetic variation lacking the ability to delineate distinct processes of evolution limit opportunities to identify determinants of fitness in rich and complex data afforded by environmental metagenomes. Indeed, the application of RSA and DTL to SAR11 demonstrate that not all variants are created equal; a notion considered common knowledge by all life scientists, and yet such a treatment is lacking in studies of genomic heterogeneity that rely upon metagenomic read recruitment. RSA and DTL provide quantitative means to bring a level of scrutiny to distinguish variants based on their distributions in proteins. For instance, a collection of high-RSA and high-DTL sites will be more likely to be enriched in neutral variants. In contrast, residues under strong purifying selection will more likely be enriched in low-RSA and/or low-DTL sites of proteins. The ability to tease apart distinct evolutionary processes with absolute accuracy will indeed remain difficult due to a multitude of factors. But by providing structure-informed means to partition the total intra-population variation into distinct pools, RSA and DTL offer a quantitative framework that enables new opportunities to study distinct evolutionary processes.

### Measuring purifying selection between genes and environments with pN/pS^(gene)^

So far, our structure-informed investigation has focused on trends of sequence variation within the gene pool of an environmental population. Next, we shifted our attention to individual proteins. pN/pS^(gene)^ is a metric that quantifies the overall direction and magnitude of selection acting on a single gene (Schloissnig et al. 2013; Shenhav and Zeevi 2020), where pN/pS^(gene)^ < 1 indicates the presence of purifying selection, the intensity of which increases as the ratio decreases. Since pN/pS^(gene)^ is defined for a given gene in a given sample, pN/pS^(gene)^ values for a single gene can be compiled from multiple samples, enabling the tracking of selective pressures across environments (Shenhav and Zeevi 2020). Taking advantage of the large number of metagenomes in which 1a.3.V was present, we calculated pN/pS^(gene)^ for all 799 protein-coding core genes across 74 samples (see Methods), resulting in 59,126 gene/sample pairs (Table S9). We validated our calculations by comparing sample-averaged pN/pS^(gene)^ to dN/dS^(gene)^ calculated from homologous gene pairs between HIMB83 and HIMB122, another SAR11 isolate genome that is closely related to HIMB83 (gANI: 82.6%), which we found to yield commensurable results (Figure S10, Table S12, Supplementary Information).

We found significantly more pN/pS^(gene)^ variation between genes of a given sample (‘gene-to-gene’ variation) than between samples of a given gene (‘sample-to-sample’ variation) (ANOVA, Figure S11). All but one gene (gene #2031, unknown function) maintained pN/pS^(gene)^ << 1 in every sample, whereby 95% of values were less than 0.15 (Figure S12, Table S9), indicating an intense purifying selection for the vast majority of 1a.3.V genes across environments. This was foreshadowed by our earlier analysis in which pS^(site)^ outweighed pN^(site)^ by 19:1 within the aggregated data across genes and samples. However, the magnitude of purifying selection was not uniform across all genes. In fact, gene-to-gene variance, as opposed to sample-to-sample variance, explained 93% of pN/pS^(gene)^ variation (ANOVA, Figure S11). By analyzing the companion metatranscriptomic data (Salazar et al. 2019) that were available for 50 of the 74 metagenomes, we were able to explain 29% of gene-to-gene variance with gene transcript abundance (Table S13, Supplementary Information), a known predictor of evolutionary rate (Pál, Papp, and Hurst 2001). Overall, these data demonstrate the utility of pN/pS^(gene)^ as a metric to understand the overall extent of selection acting on genes.

The amount of pN/pS^(gene)^ variation attributable to sample-to-sample variance was only 0.7% (Figure S11). While it represents a small proportion of the total variance, the sample-to-sample variance in pN/pS^(gene)^ encapsulates the extent that polymorphism varies in response to the range of environmental parameters observed across samples. These data therefore provide the opportunity to relate how differences in genetic diversity of individual genes manifests from differences in environmental parameters (Table S10), which we focused on next.

### Nitrogen availability governs rates of non-ideal polymorphism at critical sites of glutamine synthetase

To gain a more highly resolved picture of how selection shapes protein evolution, we searched for a biologically relevant gene within 1a.3.V that exhibited evolutionary patterns that could be understood by leveraging structural information. Glutamine synthetase (GS) is a critical enzyme for the recycling of cellular nitrogen (Bernard and Habash 2009), a limiting nutrient for microbial productivity in surface oceans (Bristow et al. 2017). GS yields glutamine and ADP from glutamate, ammonia, and ATP, an essential step in the biosynthesis of nitrogenous compounds.

Given the central role that GS plays in nitrogen metabolism, we expected GS to be under high selection. Indeed, the sample-averaged pN/pS^(GS)^ was 0.02, ranking GS amongst the top 11% most purified genes (Figure 3b, Table S9). Although highly purified, we observed significant sample-to-sample variation in pN/pS^(GS)^ (min = 0.010, max = 0.036) suggesting that the strength of purifying selection on GS varies from sample to sample (Figure 3b inset), perhaps due to unique environmental conditions (*e.g.,* nutrient compositions) that differentially impact the need for glutamine synthesis. Since previous work has shown that SAR11 upregulates its transcriptional and translational production of GS in response to nitrogen limitation (Smith Daniel P. et al., n.d.), we hypothesized that purifying selection should be highest in nitrogen-limited environments, and lowest in nitrogen-replete environments. We utilized measured concentrations of nitrate as an indication of the level of nitrogen limitation in each sample, and found a positive correlation between measured nitrate concentrations and pN/pS^(GS)^ values across samples (Pearson correlation p-value = 0.009, R^2^ = 0.11) (Figure 3c), which ranked amongst the top 12% of positive correlations between pN/pS^(gene)^ and nitrate concentration (Figure 3c inset, Table S10). In summary, we find that although GS is under high selection, subtle differences in selection strength are observed between samples and are most likely driven by nitrogen availability.

**Figure 3.**
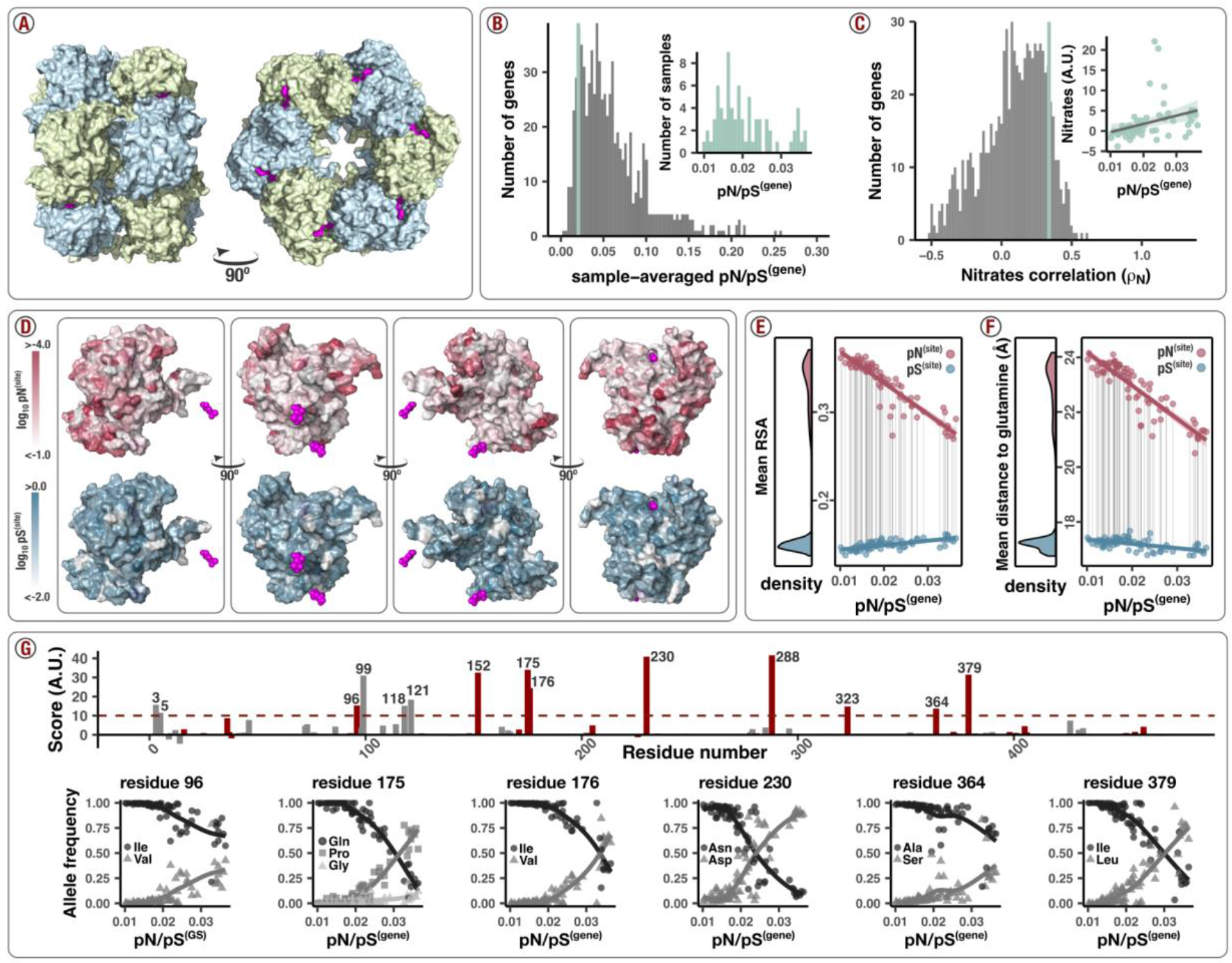
Polymorphism distribution patterns in glutamine synthetase (GS). (A) GS forms a dodecameric complex. The structure (PDB ID 1FPY) comes from *Salmonella typhimurium* (61% sequence similarity to HIMB83) and is shown from two different views. Pink molecules are ADP and phosphinothricin (steric inhibitor of glutamate), and are situated within the active site of GS. **(B) GS is one of the most highly conserved genes in 1a.3.V.** The main plot shows the distribution of sample-averaged pN/pS^(gene)^ for all 799 genes in the 1a.3.V core (truncated at 0.30). The vertical green line depicts the sample-averaged pN/pS^(gene)^ for GS (0.020). The inset plot shows the distribution of pN/pS^(gene)^ value for GS as seen across the 74 samples, which vary from 0.010 to 0.036. **(C) Selection strength on GS correlates with environmental concentration of nitrates**. The main plot shows a histogram of Pearson correlation coefficients (one per gene) between pN/pS^(gene)^ and measured concentration of nitrates in each sample. The vertical green line depicts the correlation coefficient for GS (0.34). The inset shows a scatter plot of pN/pS^(gene)^ vs nitrate concentrations from which the GS correlation coefficient was calculated. **(D) ns-polymorphism polymorphism rates are reduced in the vicinity of the active sites.** Each image is a view of the predicted structure of monomeric GS. Phosphinothricin substrates were situated by aligning the predicted GS structure to the complex in panel A. Red surfaces are colored according to the sample-averaged log10pN^(site)^ value of each residue, and blue surfaces are colored according to the sample-averaged log10pS^(site)^ value of each residue. In each case, darker colors refer to higher rates. Left-to-right, each view is a 90° clockwise rotation of the previous view about the vertical axis. Each image was rendered programmatically using a PyMOL script that was generated from the anvi’o structure interactive interface. **(E) As selection decreases, ns-polymorphism creeps into low-RSA sites.** The left panel shows the distribution of samples’ average RSA of nonsynonymous (red) and synonymous (blue) polymorphisms. The right panel shows how these average RSA values (y-axis) correlate with the samples’ pN/pS^(gene)^ values (x-axis). Each data point is calculated by weighting the RSA of each residue by the pN^(site)^ (red) or pS^(site)^ (blue) values observed in that sample. The red and blue lines show the nonsynonymous and synonymous linear fits, respectively, and the corresponding shaded regions show the 95% confidence intervals for the fit. **(F) As selection decreases, ns-polymorphism creeps closer to the binding site.** The scheme is identical to panel E, where RSA is replaced with the distance-to-glutamate substrate (DTL). **(G) Some sites exhibit amino acid minor allele frequencies that co-vary with pN/pS^(GS)^.** The top panel shows the extent that sites co-vary with pN/pS^(GS)^. The x-axis shows the residue number and the y-axis the slope estimate of a linear regression between the sum of minor allele frequencies and pN/pS^(GS)^. Sites with DTL values less than the average are indicated in red and are gray otherwise. All sites above an arbitrary cutoff (dashed horizontal line) are annotated with their residue number. Scatter plots below show the allele frequency trajectories for a select number of these sites.

Next, we focused on the GS protein structure to further investigate the associations between GS polymorphism and processes of selection. Since the native quaternary structure of GS is a dodecameric complex (12 monomers), our monomeric estimates of RSA and DTL are unrepresentative of the active state of GS. We addressed this by aligning 12 copies of the predicted structure to a solved dodecameric complex of GS in *Salmonella typhimurium* (PDB ID 1FPY), which HIMB83 GS shares 61% amino acid similarity with (Figure 3a). From this stitched quaternary structure we recalculated RSA and DTL, and as expected, this yielded lower average RSA and DTL estimates due to the presence of adjacent monomers (0.17 versus 0.24 for RSA and 17.8Å versus 21.2Å for DTL). With these quaternary estimates of RSA and DTL, we found that ns-polymorphism was 30x less common than s-polymorphism, and it strongly avoided sites with low RSA and the three glutamate active sites to which any given monomer was proximal (Figure 3d). In comparison, s-polymorphism distributed relatively homogeneously throughout the protein, whereby 17% of s-polymorphism occurred within 10Å of active sites (compared to 3% for ns-polymorphism) and 19% occurred in sites with 0 RSA (compared to 9% for ns-polymorphism).

Averaged across samples, the mean RSA was 0.15 for s-polymorphism and 0.33 for ns- polymorphism (Figure 3e left panel). Similarly, the mean DTL was 17.2Å for s-polymorphism and 22.9Å for ns-polymorphism (Figure 3f left panel). These observations highlight in a single gene what we previously observed across the 1a.3.V core: selection purifies the majority of ns- polymorphism and does so with increased strength at structurally/functionally critical sites.

We next investigated whether variance in selection strength (Figure 3b inset) affects the spatial distribution patterns of polymorphism. For each sample, we calculated how polymorphism rates in GS distributed with respect to RSA and DTL and associated these distributions with pN/pS^(GS)^. While the mean RSA of s-polymorphism remained relatively invariant (standard deviation 0.005) (Figure 3e right panel), the mean RSA of ns-polymorphism varied dramatically from 0.27 to 0.37 and was profoundly influenced by sample pN/pS^(GS)^; samples exhibiting low selection of GS harbored lower mean RSA and samples exhibiting high selection of GS harbored higher mean RSA (Figure 3e right panel). In fact, 82.9% of mean RSA ns-polymorphism variance could be explained by pN/pS^(GS)^ alone (Pearson correlation, p-value < 1x10^-16^, R^2^ = 0.829). ns- polymorphism distributions with respect to DTL were equally governed by selection strength, where 80.4% of variance could be explained by pN/pS^(GS)^ (Pearson correlation, p-value < 1x10^- 16^, R^2^ = 0.804, Figure 3f).

When selection is low, we observe high nitrate concentrations (Figure 3c inset) and ns- polymorphism distributions towards lower RSA/DTL (Figures 3e, 3f). When selection is high, we observe low environmental nitrate concentrations (Figure 3c inset) and ns-polymorphism distributions towards higher RSA/DTL (Figures 3e, 3f). Given that proper functionality of GS is most critical in nitrogen-limited environments and that mutations with low RSA/DTL are more likely to be deleterious, the most likely explanation for the body of evidence presented is that GS accumulates non-ideal polymorphism in samples exhibiting low selection of GS that cannot be effectively purified at the given selection strength. As selection increases, so too does the purifying efficiency, which we indirectly measure as increases in mean RSA and DTL of ns- polymorphism. Our approach illustrates this ‘use it or lose it’ evolutionary principle over a spectrum of selection strengths which have been sampled from natural *in situ* environmental conditions.

Under this hypothesis, there should exist low DTL amino acid alleles that create a negative, yet tolerable impact on fitness when selection is low, yet incur an increasingly detrimental fitness cost as selection increases. One would expect such alleles to be at low frequency in low pN/pS^(GS)^ samples, and to reach increasingly higher frequencies in higher pN/pS^(GS)^ samples. We identified putative sites fitting this description by scoring sites based on the extent that their amino acid minor allele frequencies co-varied with pN/pS^(GS)^, including only sites with DTL less than the mean DTL of ns-polymorphisms (22.9Å). Using an arbitrary cutoff, we identified 9 top-scoring polymorphisms that co-varied with pN/pS^(GS)^ (Figure 3g): I96V, L152I, Q175P/G, I176V, N230D, S288A/D, I323V, A364S, I379L. Though each of these sites exhibited DTL lower than the average ns-polymorphism, the closest site (residue number 323) was still 9Å away from the glutamate substrate. This suggests there are no ‘smoking gun’ polymorphisms occurring in the binding site that abrasively disrupt functionality. After all, in absolute terms GS is highly purified regardless of sample – the largest pN/pS^(GS)^ is 0.036, which is just over half the genome-wide average pN/pS^(gene)^ of 0.063. Our data therefore represents a subtle, yet resolvable signal of minute decreases in selection strength manifesting as minute shifts in the distribution of ns-polymorphism towards the active site.

While identifying signatures of positive selection is typically the primary pursuit in evolutionary analysis, our data instead illustrates a highly resolved interplay between purifying selection strength and polymorphism distribution. The geography and unique environmental parameters associated with each sample yielded a spectrum of selection strengths which enabled us to quantify how polymorphism distributions of a gene under high selection shift in response to small perturbations in selection strength. In the case of GS, we were able to attribute these shifts to the availability of nitrogen, thereby linking together environment, selection, and polymorphism.

Throughout the 1a.3.V core genes, we observed that samples exhibiting low overall selection of 1a.3.V were strongly associated with increased accumulation of ns-polymorphism at low RSA/DTL sites (Figures 4a, 4b, Supplementary Information), suggesting this signal is not specific to GS, but rather a general feature of the 1a.3.V core genes. Though highly significant (one sided Pearson p-values 9x10^-12^ for RSA and 2x10^-4^ for DTL), the magnitude that ns-polymorphism distributions shift with respect to DTL and RSA were subtle: across samples, the mean DTL of ns-polymorphism varied by less than 1Å, and the mean RSA varied between 0.230 and 0.236.

**Figure 4.**
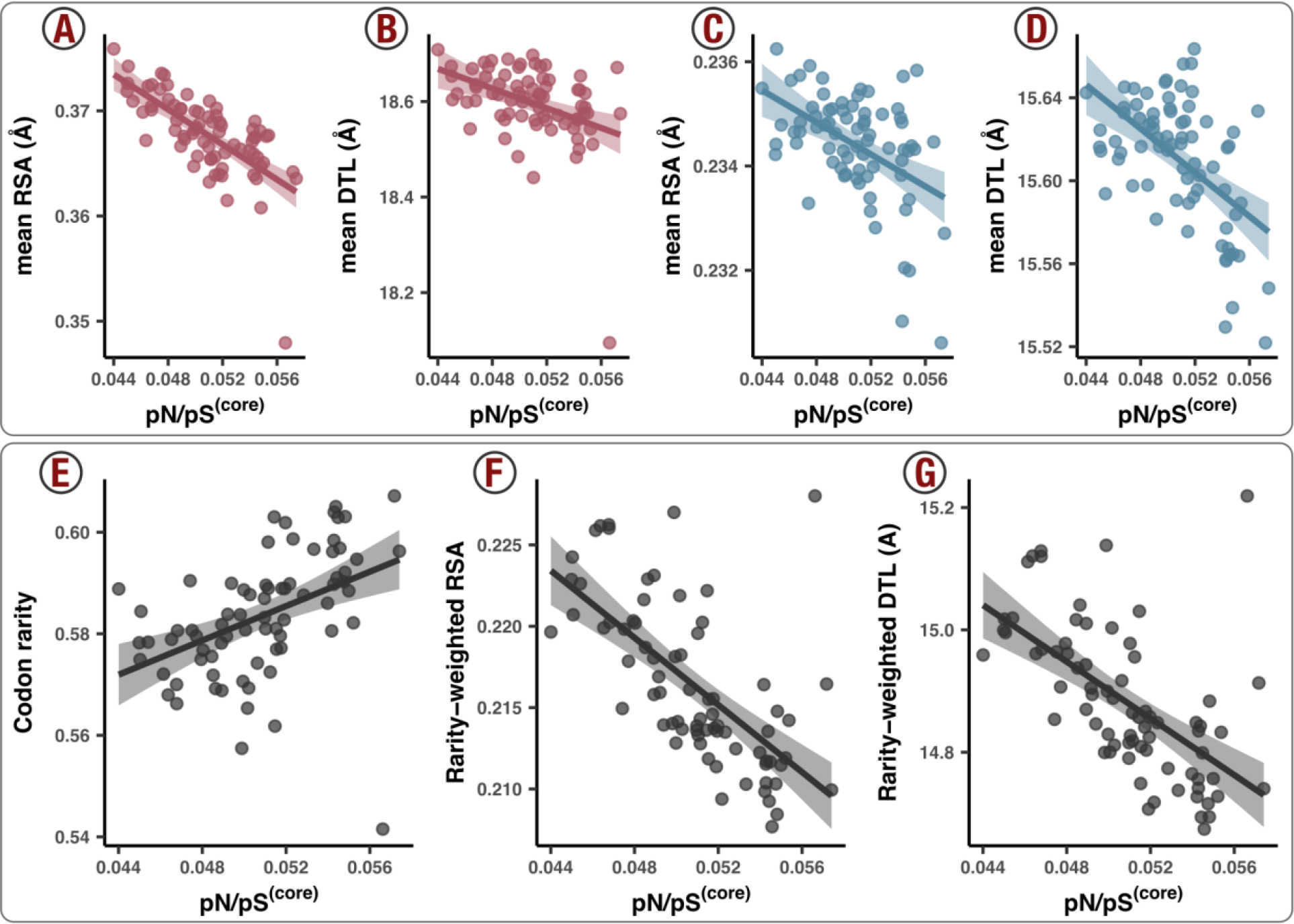
Polymorphism distribution patterns with respect to genome-wide selection strength. Each data point is a sample (metagenome). Lines represent lines of best fit and corresponding translucent areas represent 95% confidence intervals. The x-axis is pN/pS^(core)^, which is calculated across the whole core genome and is an inverse proxy of genome-wide purifying selection strength (see Methods). **(A)** The ns-polymorphism distribution mean with respect to RSA is negatively associated with pN/pS^(core)^ (one-sided Pearson p-value = 9x10^-12^). **(B)** The ns- polymorphism distribution mean with respect to DTL is negatively associated with pN/pS^(core)^ (one-sided Pearson p-value = 2x10^-4^). **(C)** The s-polymorphism distribution mean with respect to RSA is negatively associated with pN/pS^(core)^ (one-sided Pearson p-value = 1x10^-5^). **(D)** The s-polymorphism distribution mean with respect to RSA is negatively associated with pN/pS^(core)^ (one-sided Pearson p-value = 3x10^-7^). **(E)** Rare synonymous codons are more abundant in samples with high pN/pS^(core)^ (one-sided Pearson p-value = 4x10^-5^). **(F)** Rare synonymous codons avoid low RSA sites when pN/pS^(core)^ is low (one-sided Pearson p-value = 1x10^-10^). **(G)** Rare synonymous codons avoid low DTL sites when pN/pS^(core)^ is low (one-sided Pearson p-value = 7x10^-9^).

Resolving such a minute signal with such robust statistical power is owed to the immense quantities of sequence data afforded by metagenomics.

### Synonymous but not silent: selection against rare codons at critical sites

Thus far we have observed that purifying efficiency observably decreases in response to lowered selection strength, as evidenced by ns-polymorphism occurring nearer to binding sites and in more buried sites. Given the influence of synonymous substitutions in translational processes (Plotkin and Kudla 2011), as a final analysis we focused on within-population trends of s- polymorphism.

Compared to ns-polymorphism, s-polymorphism distributes more uniformly throughout protein structures (Figures 2a, 2b). Yet our data also revealed an association between selection strength and the distribution of s-polymorphism. In samples under higher selection, s-polymorphism systematically tended to occur (1) in more solvent-exposed sites (Figure 4c, one-sided Pearson p-value = 1x10^-5^) and (2) farther from binding sites (Figure 4d, one-sided Pearson p-value = 3x10^- 7^). These trends indeed mimic the nonsynonymous trends in glutamine synthetase (Figures 3d, 3e, 3f) as well as the core genes in general (Figures 4a, 4b), and cannot be reasonably explained by neutral processes. The surprising association suggests a relationship between selection and synonymous change that is at least partly determined by structural features of proteins.

With a GC-content lower than 30%, SAR11 genomes maintain a non-uniform yet conserved codon composition (Figure S13). Previous work has shown that rare codons can significantly reduce translation rates (Sørensen, Kurland, and Pedersen 1989), cause delays in the production of the polypeptide chain at the ribosome (Komar 2009), which can lead to protein misfolding (Drummond and Wilke 2008; Agashe et al. 2013), and impair fitness (Walsh et al. 2020). Thus, we hypothesized that rare codons in 1a.3.V may incur fitness costs relative to their more common, synonymous counterparts. To test this hypothesis, we investigated the relationship between selection strength and the occurrence of rare codons, which required us to define a ’codon rarity’ metric based on the frequency that codons are found in the HIMB83 genome relative to their synonymous counterparts (Table S11). We then attributed an overall rarity score to each sample by weighting the rarity of all synonymous codon alleles by the frequencies with which they were observed (see Methods). Our analysis of these data revealed a positive correlation between codon rarity in a sample and its pN/pS^(core)^ (Figure 4d, one-sided Pearson p-value = 1x10^-5^), illustrating that rare codons are more likely to be found in samples where genome-wide selection is low. We found this to be the case for s-polymorphism within all 18 amino acids that possess two or more codons (Figure S14), illustrating that this evolutionary process acts ubiquitously throughout the genetic code of 1a.3.V. Rare codons did not distribute throughout protein structures uniformly, either. In samples with low genome-wide selection, where rarity was highest, rare codons occurred farther away from binding sites (one-sided Pearson p-value = 1x10^-10^) and occurred more frequently in more solvent-exposed sites (one-sided Pearson p-value = 7x10^-9^), as compared to low selection samples (Figures 4e, 4f).

Overall, these data show that when genome-wide selection strength is low, rare codons both (1) incorporate into the genome with increased propensity, and (2) manifest in sites that are statistically more likely to be structurally/functionally important. As previous research suggests, the most likely explanation for these observations is that rare codons are less fit due to decreased translational accuracy compared to their more common, synonymous counterparts. Yet the environmental and structural dimensions of our data reveal the dynamic nature of the evolutionary processes that maintain synonymous polymorphism as a function of changing conditions in naturally occurring habitats and elucidates the intensity of such processes as a function of their physical locations in the structure. Indeed, 1a.3.V maintains the lowest proportion of rare codons in samples where genome-wide selection is highest, and rare codons in these samples are statistically more likely to be incorporated in noncritical sites of proteins, most likely due to the increased efficiency with which purifying selection operates in an environment- and site-dependent manner. These rare codon data provide a lens into the potential fitness costs associated with suboptimal translational accuracy in complex populations, and by including structural data, we demonstrate where optimal translational accuracy matters most.

## Conclusions

With recent breakthroughs in predicting protein structures and ligand binding sites, microbial ecology need not be limited to just sequences. By offering an interactive, scalable, and open-source software solution that integrates environmental genetic variants with structural bioinformatics, our study takes advantage of recent advances to connect environmental ‘omics and structural biology. Indeed, by leveraging structure and ligand-binding predictions we were able to describe striking patterns of nucleotide polymorphism in an environmental microbial population that we could ascribe to evolutionary constraints that preserve protein structure (folding & stability) and protein function (ligand-binding activity). By tracking a SAR11 population across metagenomes we were able to demonstrate the presence of dynamic processes that purge both synonymous and nonsynonymous polymorphism from the vicinity of ligand binding sites of proteins as a function of selection strength. Overall, our study proposes a structure-informed computational framework for microbial population genetics and offers a glimpse into the emerging interdisciplinary opportunities made available at the intersection of ecology, evolution, and structural biology.

## Methods

### Overview

The URL https://merenlab.org/data/anvio-structure/ provides a complete reproducible workflow for all analysis steps detailed below, including (1) downloading the publicly available metagenomes and genomes, (2) recruiting reads from metagenomes, (3) calculating single amino-acid and single codon variants, (4) predicting protein structures and ligand binding sites, and (5) visualizing metagenomic sequence variants and binding sites onto protein structures.

### Metagenomic and metatranscriptomic read recruitment and processing

To study the population structure of the environmental SAR11 population 1a.3.V defined previously (Delmont and Kiefl et al. 2019), we used anvi’o v7.1 (Eren et al. 2021), and its metagenomics workflow (Shaiber et al. 2020) which uses snakemake v5.10 (Köster and Rahmann 2012) to automate gene calling, gene function annotation, metagenomic and metatranscriptomic read recruitment steps. The compendium of anvi’o programs the metagenomics workflow called upon employed Prodigal v2.6.3 (Hyatt et al. 2010) for gene calling, NCBI’s Clusters of Orthologous Groups (COGs) database (Tatusov et al. 2003) and Pfams (El-Gebali et al. 2019) for gene function annotation, HMMER v3.3 (Eddy 2011) for profile HMM searches, DIAMOND v2.0.6 (Buchfink, Xie, and Huson 2015) for sequence searches, Bowtie2 v2.4 (Langmead and Salzberg 2012) for read recruitment, and samtools v1.9 (Li et al. 2009) to generate BAM files. The metagenomic workflow resulted in a ‘contigs database’ and a ‘merged profile database’ (two anvi’o artifacts detailed at https://anvio.org/help/), which gives access to gene and genome coverages (with metagenomic or metatranscriptomic short reads), as well as the sequence variability data to study population genetics as detailed below. We adopted a competitive read recruitment strategy by using all SAR11 genomes, rather than only HIMB83, as reference to recruit reads from Tara Oceans Project metagenomes and metatranscriptomes to maximize the exclusion of reads that matched better to other known SAR11 genomes, thereby narrowing our scope of probed diversity and minimizing the impacts of non-specific read recruitment. In all subsequent analyses we focused on the core genes of the 1a.3.V subclade by only considering (a) reads that mapped to HIMB83 (b) the 74 metagenomes in which HIMB83 was found above 50X, and (c) the 799 HIMB83 genes that were previously found to maintain consistent coverage patterns (Delmont and Kiefl et al. 2019).

### Quantifying SCVs and SAAVs in metagenomes

To characterize the variants in metagenomic read recruitment results we used and extended the microbial population genetics framework implemented in anvi’o. The program ‘anvi-profilè with the flag ‘--profile-SCVs’ characterizes single codon variants (SCVs), from which single amino acid variants (SAAVs) can also be calculated. Anvi’o determines allele frequency vectors for SCVs by tallying the frequencies of codons observed in the 3-nt segments of reads that fully map to a given codon position. The frequencies of amino acids encoded by each 3-nt segment yield SAAVs observed in a given position, which represent allele frequency vectors of positions after collapsing synonymous redundancy. For a given codon position, anvi’o excludes any reads that do not map to all 3 nucleotides, which can happen either if the read terminates within the codon position, or there exists a deletion in the read relative to the reference genome. Reads that contain insertions within the codon relative to the reference genome are also excluded during this step. We exported variant profiles as tabular data using the program ‘anvi-gen-variability-profilè, where each row is a SCV (or SAAV) and the columns specify (1) identifying information such as the corresponding gene, codon position, and sample id, (2) the number of mapped reads corresponding to each of the 64 codons (or 20 amino acids), and (3) numerous miscellaneous statistics, all of which can be explored at https://merenlab.org/analyzing-genetic-varaibility/.

### Calculations of polymorphism rates of individual codon sites, pN^(site)^ and pS^(site)^

We calculated the polymorphism rates of individual codon sites from allele frequencies defined from each SCV based on a recent study by Shenhav and Zeevi (2020), where a given codon allele contributes (to either pN^(site)^ or pS^(site)^) an amount that is equal to its observed relative abundance (frequency). To which rate the allele contributes is determined by its synonymity relative to the popular consensus, i.e. the allele most common across all samples. After summing the contributions for each of the 63 codons (excluding the popular consensus), we normalized the resulting values of pN^(site)^ and pS^(site)^ by the number of nonsynonymous and synonymous sites of the popular consensus, respectively. For example, if the popular consensus is ‘ACC’ (Thr), there are 9 possible single point mutations, 3 synonymous and 6 nonsynonymous, therefore pS^(site)^ will be divided by 3/3 = 1 and pN^(site)^ will be divided by 6/3 = 2. This procedure can be mathematically expressed as

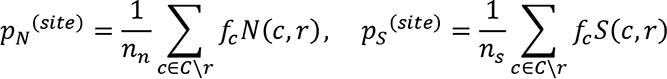

Where *C*\*r* is the set of all codons excluding the popular consensus *r*; *n*_*n*_ and *n*_*s*_ are the number of nonsynonymous and synonymous sites of *r*, respectively; *N*_*c,r*_ is the frequency of the *c*th allele;
*N*(*c*, *r*) is the indicator function where,

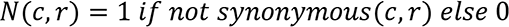

and *S*(*c*, *r*) is the indicator function where,

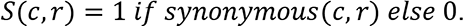

We implemented this strategy into the program ‘anvi-gen-variability-profilè as a new flag ‘-- include-site-pnps’, which when declared, adds pN^(site)^ and pS^(site)^ values as additional columns to the tabular output after calculating them for 3 different choices of the reference codon *r*: (1) the popular consensus (as used in this paper), (2) the consensus (the allele with the highest frequency), and (3) the codon found in the reference sequence (the sequence used for read recruitment). For efficient computation, this calculation uses the Python package numba (Lam, Pitrou, and Seibert 2015) for just-in-time compilation. For a dataset with 12,583,626 SCVs, the current implementation computes pN^(site)^ and pS^(site)^ terms in less than a minute on a laptop computer.

### Calculations of polymorphism rates within a group of sites, pN^(group)^, pS^(group)^, and pN/pS^(group)^

We defined groups such that all sites in a group share similar RSA and DTL values. Formally, we defined pN^(group)^ and pS^(group)^ as

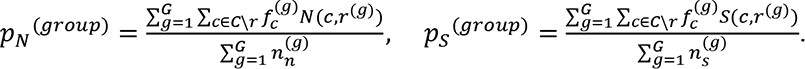

*G* is the number of sites in the group; *r*^(*g*)^ is the popular consensus of the *g*th site; *n*_*n*_ ^(*g*)^ is the frequency of the *c*th allele at the *g*th site; *g*_*r*_^(*g*)^ and *n*_*s*_^(*g*)^ are the number of nonsynonymous and synonymous sites of *r*^(*g*)^, respectively. All other definitions are the same as for pN^(site)^ and pS^(site)^. pN^(group)^ and pS^(group)^ can be expressed in terms of weighted sums of pN^(site)^ and pS^(site)^, respectively:

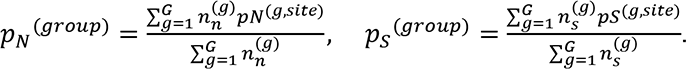

Finally, pN/pS^(group)^ is defined as

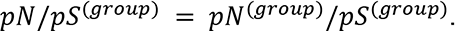

### Calculations of polymorphism rates for individual and core genes, pN^(gene)^, pS^(gene)^,pN/pS^(gene)^, and pN/pS^(core)^

We calculated rates of polymorphism for genes and the 1a.3.V core genome identically to the calculations of pN^(group)^, pS^(group)^, and pN/pS^(group)^. For example, pN^(gene)^ refers to the ns-polymorphism rate of all sites in a given gene, and pS^(core)^ refers to the s- polymorphism rate of all sites in the 1a.3.V core genome.

### Predicting and processing protein structures

We attempted to predict protein structures for each gene in the HIMB83 genome that belonged to the 1a.3.V core using both AlphaFold (Jumper et al. 2021) and MODELLER (Webb and Sali 2016). To process, store, and access the resulting protein structures we developed a novel program, ‘anvi-gen-structure-databasè, which gives access to all atomic coordinates as well as per-residue statistics such as relative solvent accessibility, secondary structure, and phi & psi angles calculated using DSSP (Touw et al., 2015; Kabsch and Sander, 1983). For AlphaFold predictions we used a version of the codebase that closely resembles v2.0.1 (<u>this URL</u> gives access to its exact state) and ran predictions using 6 GPUs, which took a week on a high-performance computing system. AlphaFold predicted structures for 795 of 799 proteins, and after removing structures with gene-averaged pLDDT scores <80, we were left with 754 structures we deemed ‘trustworthy’ for downstream analyses. To predict protein structures with MODELLER, we developed a pipeline that, for each gene, (1) searches the Research Collaboratory for Structural Bioinformatics Protein Data Bank (Berman et al. 2000) (RSCB PDB) for homologs using DIAMOND (Buchfink, Xie, and Huson 2015), then downloads tertiary structures for matching entries, and (2) uses these homologs as templates to predict the gene’s structure with MODELLER (Webb and Sali 2016). We discarded any proteins if the best template had a percent similarity of <30%. Unlike more sophisticated homology approaches that make use of multi-domain templates (Källberg et al. 2012), we used single-domain templates which are convenient and are accurate up to several angstroms, yet can lead to physically inaccurate models when the templates’ domains match to some, but not all of the sequences’ domains. To avoid this, we discarded any templates if the alignment coverage of the protein sequence to the template was <80%. Applying these filters resulted in 408 structures from the 1a.3.V core, which was further refined by requiring that the root mean squared distance (RMSD) between the predicted structure and the most similar template did not exceed 7.5 Å, and that the GA341 model score exceeded 0.95. After applying these constraints, we were left with 348 structures in the 1a.3.V that we assumed to be ‘trustworthy’ structures as predicted by MODELLER. These structures were on average 44.8% identical to their templates, which is within the sequence similarity regime where template-based homology modeling generally produces the correct overall fold (Rost 1999).

### Predicting ligand-binding sites

For the 1a.3.V core genes we estimated per-residue binding frequencies for a diverse collection of ligands by using InteracDome, a database that annotates the sites (match states) of Pfam profile hidden Markov models (HMMs) with ligand binding frequencies predicted from experimentally-determined structural data (Kobren and Singh 2019). To associate match state binding frequencies of the profile HMMs to the sites of HIMB83 genes, we applied a protocol similar to that described in Kobren & Singh.

First, we downloaded the Representable-NR Interactions (RNRI) from the InteracDome web server (https://interacdome.princeton.edu/) that “correspond to domain-ligand interactions that had nonredundant instances across three or more distinct PDB structures” (Table S5). Next, we downloaded the profile HMMs for Pfam v31.0 and kept only those 2,375 profiles that belonged to the RNRI dataset. Then, we searched each HIMB83 gene against this set using HMMER’s hmmsearch. After the removal of HMM hits that were below the gathering threshold (GA) noise cutoffs defined in Pfam models, 940 of the 1,470 HIMB83 coding genes had at least one domain hit, with a total of 1,770 domain hits from 832 unique profile HMMs. Of these, we removed 177 for being too partial (length of the hit divided by the profile HMM length was less than 0.5), and 1 hit because the query sequence did not match all the consensus residues for match states in which the information content exceeded 4 (Table S5). We then associated binding frequencies for a collection of ligand types to the HIMB83 genes by parsing alignments of the profile HMMs to the HIMB83 gene amino acid sequences, which are provided in the standard output of hmmsearch. If a given HIMB83 residue aligned to multiple match states, each which had the same ligand type, we attributed the average binding frequency to the HIMB83 residue. We then filtered out binding frequency scores less than 0.5, yielding 40,219 predicted ligand-residue interactions across 11,480 unique sites (Table S5). We considered each of these sites to be ‘ligand-binding sites’.

Our study includes two novel programs to automate this procedure and make it accessible to the community. The first, ‘anvi-setup-interacdomè, downloads the RNRI and Pfam datasets, and only needs to be run once. The second, ‘anvi-run-interacdomè, is a multi-threaded program that takes an anvi’o contigs database as input, and runs the remainder of the workflow described for each gene in the database. Predicted binding frequencies are stored internally in the database, which enables a seamless integration with other anvi’o programs to accomplish various tasks, such as the interactive visualization of the binding sites of predicted structures for any given gene with ‘anvi-display-structurè (see Supplementary Information), or exporting the underlying data as TAB-delimited files with ‘anvi-export-misc-datà. In the present study, ‘anvi-run-interacdomè processed the HIMB83 genome in 53 seconds on a laptop computer using a single thread.

### Calculating relative solvent accessibility (RSA)

We calculated RSA for each residue of each predicted structure, where RSA was defined as the accessible surface area (ASA) probed by a 1.4Å radius sphere, divided by the maximum ASA, *i.e.* the ASA of a Gly-X-Gly tripeptide. RSA values were calculated in the program ‘anvi-gen-structure-databasè using Biopython’s DSSP module (Cock et al. 2009).

### Calculating distance-to-ligand (DTL)

DTL was calculated for all sites that belonged to genes with (a) a predicted structure and (b) at least one predicted ligand-binding residue. Ideally, one would calculate DTL as the Euclidean distance of a residue to the predicted ligand, however our predictions did not yield the 3D coordinates of ligands. Instead, we approximated DTL as the Euclidean distance of a residue to the closest ligand-binding residue (see Methods), which lies within a few angstroms of the predicted ligand. Specifically, we defined this distance according to the sites’ side chain center of masses. A consequence of approximating DTL with respect to the closest ligand-binding sites is that by definition, any ligand-binding residue has a DTL of 0.

As discussed in *Proteomic trends in purifying selection are explained by RSA and DTL*, missed binding sites lead to erroneously high DTL values. We assessed the magnitude of this error source by comparing our distribution of predicted DTL values in the 1a.3.V core to that found in BioLiP, an extensive database of semi-manually curated ligand-protein complexes (Yang, Roy, and Zhang 2013). We found the 1a.3.V DTL distribution had a much higher proportion of values >40 Å, suggesting these likely result from incomplete characterization of binding sites (Figure S9). To mitigate the influence of this inevitable error source, we conservatively excluded DTL values >40 Å (8.0% of sites) in all analyses after Figure 2b.

### Calculating polymorphism null distributions for RSA and DTL

The null distributions for polymorphism rates with respect to RSA and DTL were calculated by randomly shuffling the RSA and DTL values calculated for each site, yielding distributions one would expect if there was no association between polymorphism rate and RSA. To avoid biases, each null distribution is the average of 10 shuffled datasets.

### Proportion of polymorphism rate variance explained by RSA and DTL

To calculate the extent that RSA and DTL can explain polymorphism rates, we constructed 3 synonymous models (s-models) and 3 nonsynonymous models (ns-models) (Table S6). s-models fit linear regressions of log10(pS^(site)^) to RSA (s #1), DTL (s #2), and both RSA & DTL (s #3). Similarly, ns-models fit linear regressions of log10(pN^(site)^) to RSA (ns #1), DTL (ns #2), and both RSA & DTL (ns #3). Additionally, each model included the gene and sample of the corresponding polymorphism as independent variables, in order to account for gene-to-gene and sample-to-sample differences. Polymorphism rates were log-transformed because it helped linearize the data, yielding better models. The data used to fit each model included all codon positions across all samples in each gene that had a predicted protein structure and at least 1 predicted ligand-binding residue. After excluding monomorphic sites (pN^(site)^ = 0 for ns-models, pS^(site)^ = 0 for s-models), this yielded 5,838,445 data points for s-models and 3,850,182 for ns-models. While every protein has RSA values that span the domain [0,1], protein size creates dramatic gene-to-gene differences in observed DTL values. We accounted for this by standardizing DTL values on a per-gene basis, which improved variance explained by DTL. The variance explained by RSA, DTL, sample, and gene was determined by performing an ANOVA on each model and partitioning the sum of squares (Table S6).

### Calculating transcript abundance (TA)

Since proper transcription level metrics such as molecules per cell are incalculable from metatranscriptomic data, we estimated the transcript abundance (TA) to be

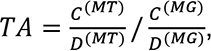

Where *C*^(*MT*)^ is the coverage of the gene in the metatranscriptome, *D*^(*MT*)^ is the sequencing depth (total number of reads) of the metatranscriptome, *C*^(*MG*)^ is the coverage of the gene in the metagenome, and *D*^(*MT*)^ is the sequencing depth (total number of reads) of the metagenome.

This means, for example, that a gene with a metatranscriptomic relative abundance 10% of its metagenomic relative abundance would have a TA of 0.10.

### Definition of codon rarity

We defined the rarity of a codon in the following way. First, we calculated the unnormalized codon rarity for each codon *c*, which we defined as

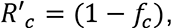

where *f*_*c*_ is the frequency that a codon was observed in the HIMB83 genome sequence. Then, we normalized the values such that the codons with the lowest and highest values get rarity scores of 0 and 1, respectively:

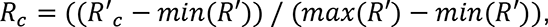

where *min*(*R*′) and *max*(*R*′) correspond to the smallest and largest unnormalized rarity scores.

We utilized this definition to calculate codon rarity at polymorphic sites by weighting each codon’s rarity by the frequency that the codon was observed in the short reads mapping to that position. For example, a polymorphic site with a coverage of 200, where 50 reads resolve to GCC (*R*_*GCC*_ = 0.97) and 150 resolve to GCT (*R*_*GCC*_ = 0.75) would get a rarity score of 50/200 × 0.97 + 150/200 × 0.75 = 0.81. Extending this to multiple sites, we take the codon rarity of an entire sample to be the average rarity across all codon sites (polymorphic or not).

### Statistical data analysis and visualization

We used R v3.5.1 (R Development Core Team 2011) for the analysis of numerical data reported from anvi’o. For data visualization we used ggplot2 (Ginestet 2011) library in R and anvi’o, and finalized images for publication using Inkscape v1.1 (https://inkscape.org/).

## Acknowledgements

We thank Stephen Giovannoni (https://microbiology.oregonstate.edu/dr-stephen-giovannoni), Shilpa Nadimpalli Kobren (http://shilpakobren.com/), Michael K. Yu (@michaelkuyu), Tobin Sosnick (http://sosnick.uchicago.edu/), as well as the members of our laboratory (https://merenlab.org/people/) for helpful discussions. EK acknowledges support from the Natural Sciences and Engineering Research Council of Canada. This work was supported by Alfred P. Sloan Foundation Fellowship in Ocean Sciences to AME, and by a Simons Foundation grant (#687269) to AME.

## Author contributions

EK and AME conceptualized the study and interpreted findings. EK curated data, developed software tools, and performed primary analyses. OCE, and AME contributed software. EK and AME wrote the paper. SEM, KK, and ADW helped with data analyses and interpretation. MSP and TP helped with project management and funding acquisition. AME supervised the project. All authors commented on the drafts of the study. All authors read and approved the final manuscript.

## Supplementary Tables

Table S1. Read recruitment and coverage statistics of the 21 SAR11 genomes. **(A-D)** Genome-wide statistics for each genome in each metatranscriptomic and metagenomic sample. **(A)** is the mean coverage, **(B)** is the mean coverage, excluding nucleotide coverage values outside the interquartile range (IQR), **(C)** is the detection, and **(D)** is the percentage of reads mapping to a genome (sums to 100 for a given sample) **(E)** The mean coverage of each HIMB083 gene in each metatranscriptomic and metagenomic sample.

Table S2. Average percent similarity of recruited reads by HIMB083 for each **(A)** gene-sample pair, **(B)** gene (marginalized over samples), and **(C)** sample (marginalized over genes).

Table S3. Mean per-site polymorphism rates (pN^(site)^ and pS^(site)^) of HIMB083 **(A)** over all sites, genes, and samples, as well as **(B)** for each gene-sample pair **(C)** each gene (marginalized over samples), and **(D)** each sample (marginalized over genes).

Table S4. Methodological comparisons between AlphaFold and MODELLER structures. **(A)** Key metrics for AlphaFold- and MODELLER-predicted structures and their alignments. **(B)** PDB structures used as templates for MODELLER predictions. **(C)** Per-residue pLDDT scores for AlphaFold-predicted structures. **(D)** Gene-averaged pLDDT scores for AlphaFold-predicted structures. **(E-F)** Genes with AlphaFold and MODELLER structures, respectively, that we determined to be of sufficiently high quality.

**Table S5.** Summary of ligand-binding residue predictions with InteracDome. **(A)** All predicted ligand-binding sites, the predicted ligand, and the predicted ligand binding score. **(B)** Characterization of each HMM domain hit. **(C)** Each match state from the Pfam profile HMMs that contributed to each predicted ligand-binding residue of HIMB083.

|  |  | Sum of squares (% of total) |  |  |  |  |
| --- | --- | --- | --- | --- | --- | --- |
| Name | Model | RSA | DTL | Gene | Sample | Residuals |
| s #1 | $\log_{10}(pS^{(site)}) \sim \text{RSA} + \text{gene} + \text{sample}$ | 0.12 | - | 4.05 | 1.01 | 94.83 |
| ns #1 | $\log_{10}(pN^{(site)}) \sim \text{RSA} + \text{gene} + \text{sample}$ | 11.83 | - | 10.61 | 6.71 | 70.85 |
| s #2 | $\log_{10}(pS^{(site)}) \sim \text{DTL} + \text{gene} + \text{sample}$ | - | 0.30 | 4.07 | 1.00 | 94.63 |
| ns #2 | $\log_{10}(pS^{(site)}) \sim \text{DTL} + \text{gene} + \text{sample}$ | - | 6.89 | 12.08 | 7.10 | 74.39 |
| s #3 | $\log_{10}(pS^{(site)}) \sim \text{RSA} + \text{DTL} + \text{gene} + \text{sample}$ | 0.03 | 0.30 | 4.05 | 1.00 | 94.62 |
| ns #3 | $\log_{10}(pS^{(site)}) \sim \text{RSA} + \text{DTL} + \text{gene} + \text{sample}$ | 7.23 | 6.89 | 10.83 | 6.42 | 68.62 |

Table S6. Summary of models used for estimating the explanatory power of RSA and DTL on polymorphism rates (see Methods).

Table S7. Summary statistics for the polymorphism models of gene-sample pairs.

Table S8. Summary of per-group polymorphism data for **(A)** pN^(group)^, **(B)** pS^(group)^, **(C)** pN/pS^(group)^, and **(D)** the size of each group.

Table S9. Summary of per-gene polymorphism data for **(A)** pN/pS^(gene)^, **(B)** sample-averaged pN/pS^(gene)^, **(C)** pN^(gene)^, **(D)** pS^(gene)^ and **(E)** the number of potential synonymous and nonsynonymous point mutations of each gene.

Table S10. Correlations of pN/pS^(gene)^ for each 1a.3.V core gene with respect to the measured environmental parameters: nitrates, chlorophyll, temperature, salinity, phosphate, silicon, depth, and oxygen.

Table S11. Codon metrics, including anti-codon, encoded amino acid, frequency and rarity in genome, and frequency and rarity compared to synonymous codons.

Table S12. Comparison between dN/dS between HIMB83 and HIMB122 homologs and sample-averaged pN/pS^(gene)^ of 1a.3.V genes.

Table S13. Per sample and gene measures of transcript abundance (TA) and related quantities.

Table S14. Bootstrap estimates of Pearson correlation coefficients and p-values from Figure SI6.

## Supplementary Figures

**Figure S1.**
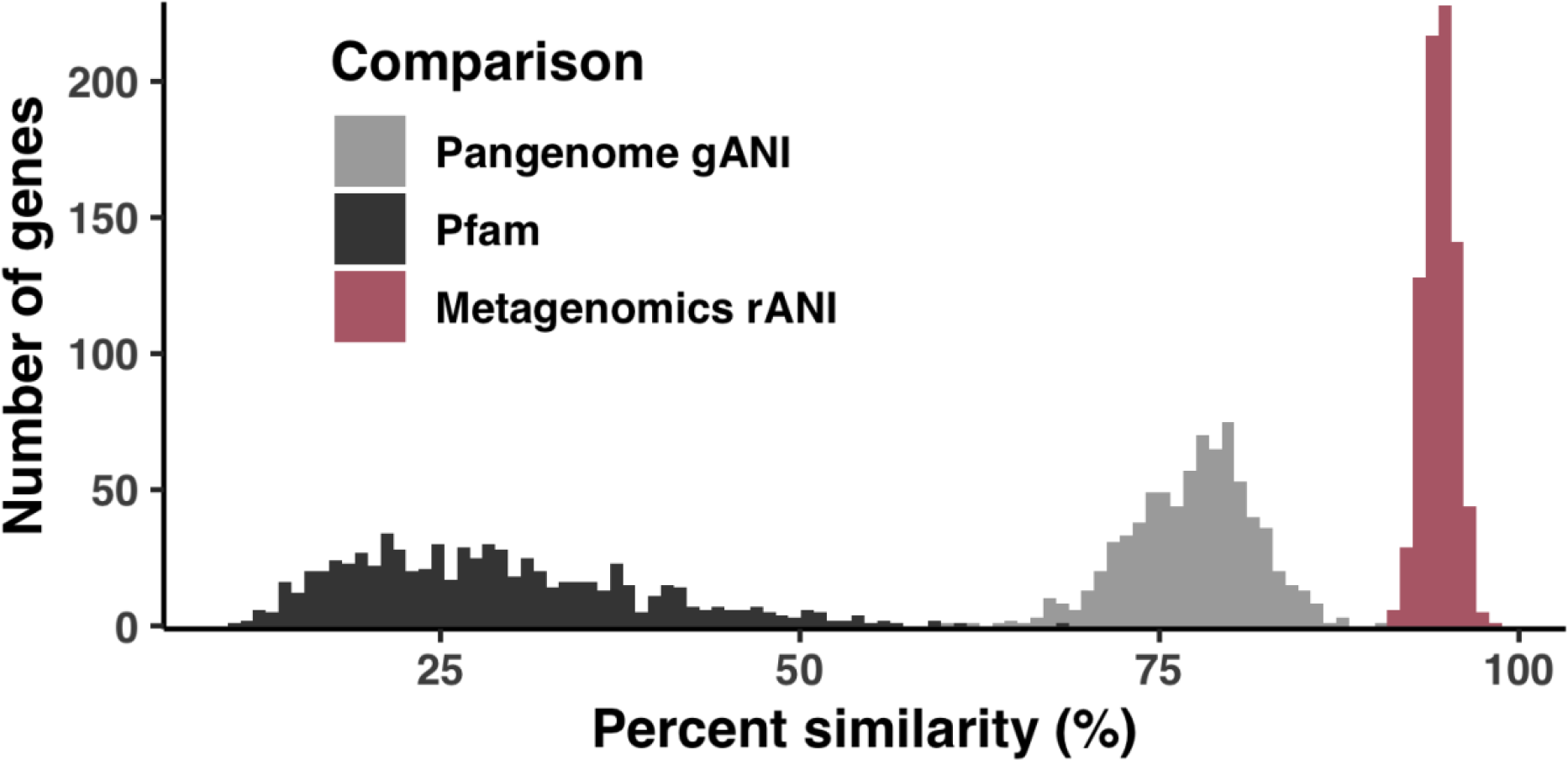
Regimes of sequence similarity probed by metagenomics, SAR11 cultured genomes, and protein families. Empirical distributions of gene-level percent similarity for HIMB83 compared with recruited metagenomic reads (pink), homologous SAR11 genomes (blue), and homologous Pfams (orange). For calculation details, see Supplementary Information.

**Figure S2.**
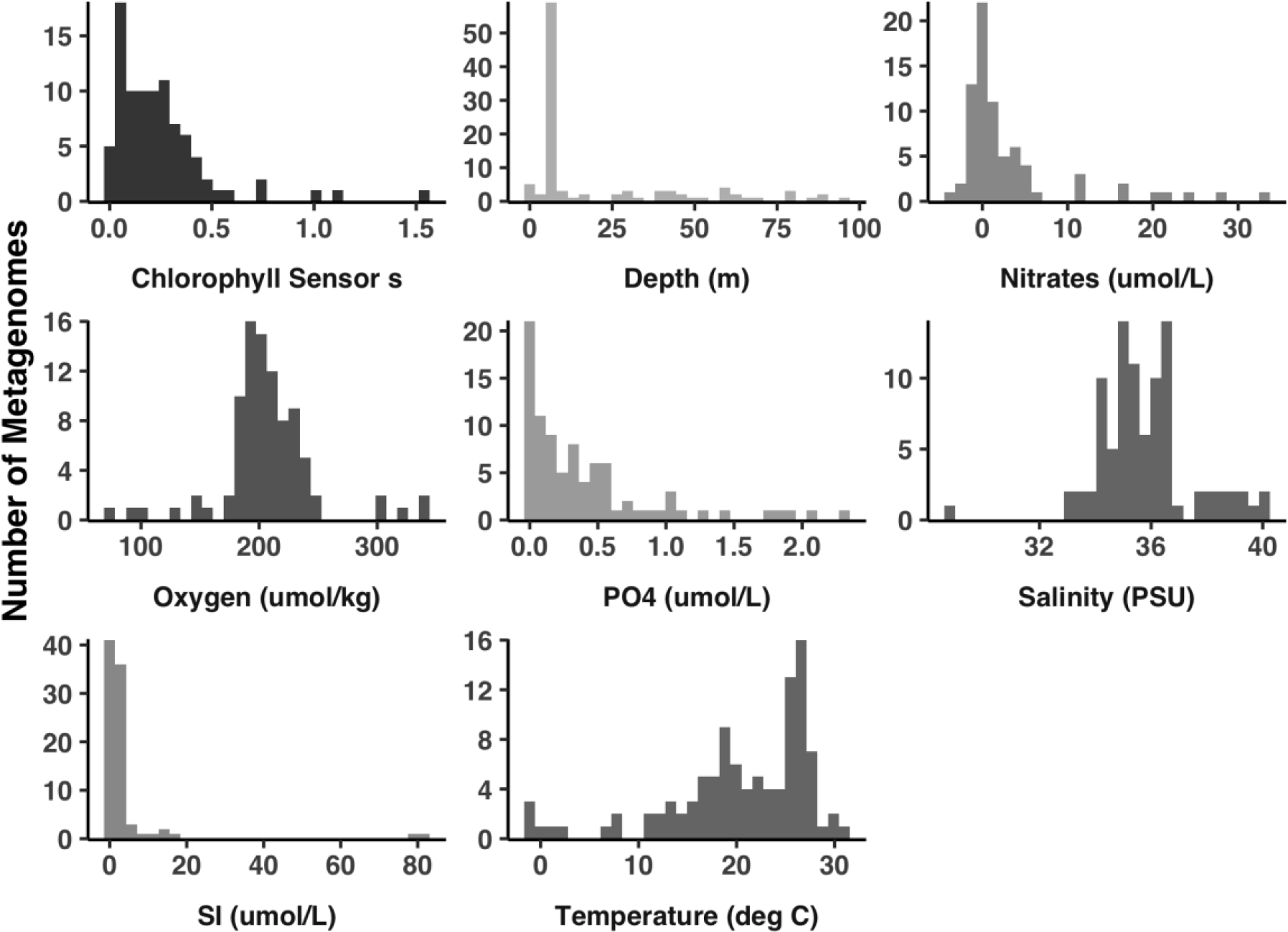
Different environments exhibit substantial variation in their environmental parameters. Each subplot shows how the 74 selected metagenomes distribute according to various environmental variables measured by the TARA ocean metagenome project.

**Figure S3.**
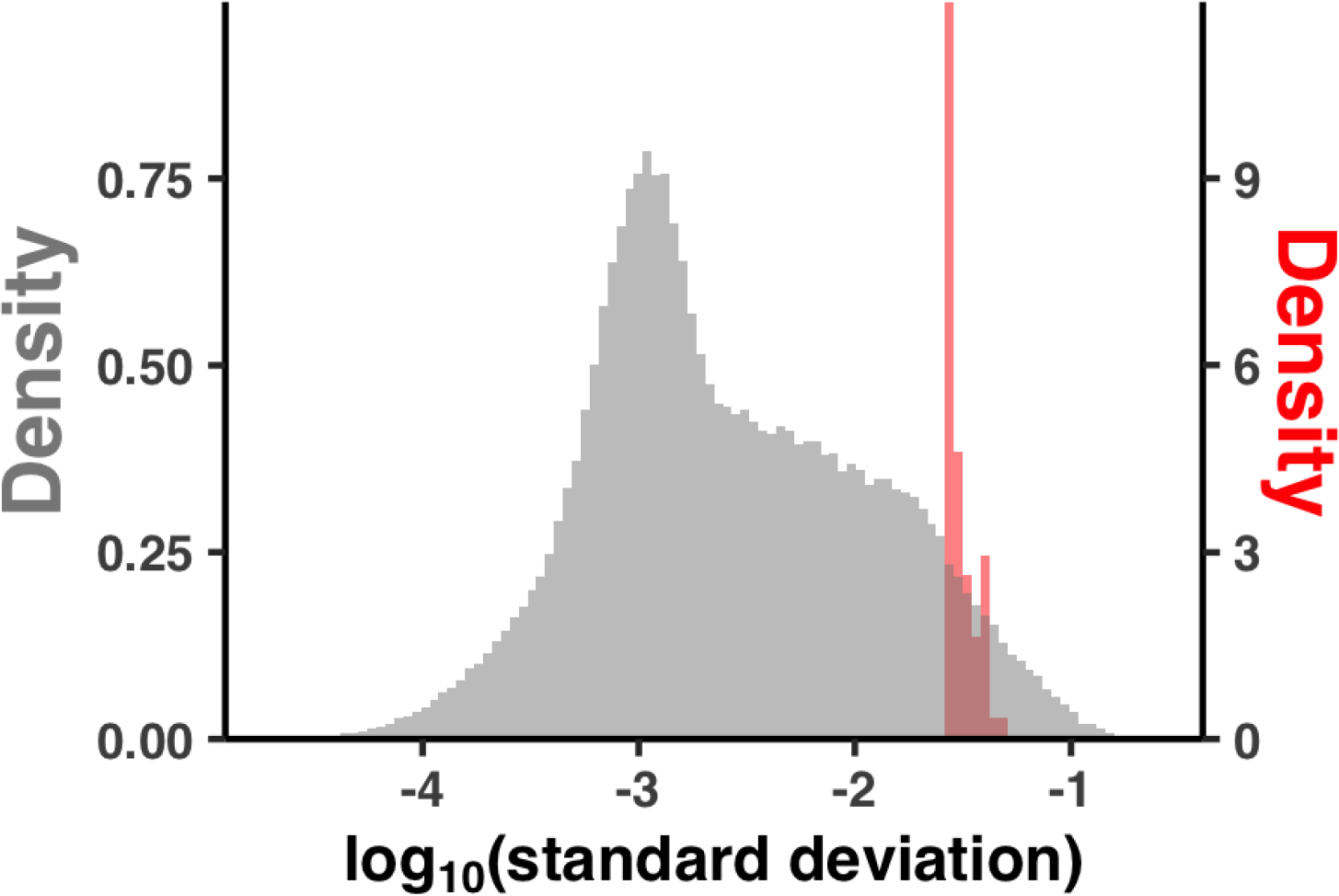
pN^(site)^ varies more significantly between sites in a given sample than between samples for a given site. The x-axis is the log-transformed standard deviation of either a sample’s pN^(site)^ values observed over many sites (orange), or a site’s pN^(site)^ values observed over the 74 samples (gray).

**Figure S4.**
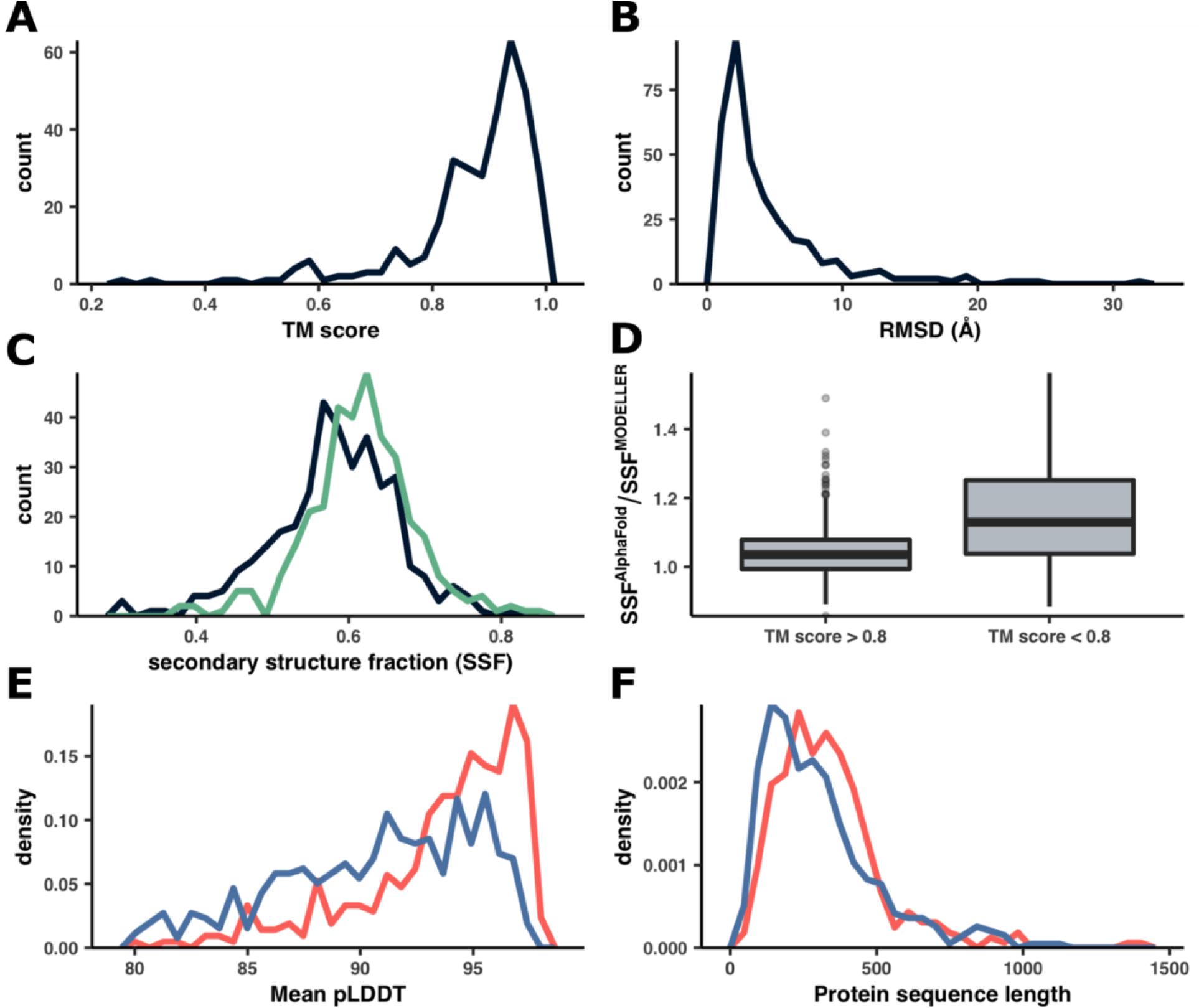
Comparisons between structures predicted by AlphaFold and MODELLER. (A-B) Distributions of TM scores and RMSD between structures predicted by both MODELLER and AlphaFold. **(C)** Distribution of secondary structure fractions, between MODELLER (black) and AlphaFold (green). Secondary structure fraction was defined for each gene as the fraction of sites that DSSP predicted as part of an alpha helix or beta strand. **(D)** Comparison of secondary structure fractions between MODELLER and AlphaFold for two TM score groups. The y-axis is the secondary structure fraction of AlphaFold divided by the secondary structure fraction of MODELLER. The two groups were defined as having TM scores above or below 0.8, where the >0.8 group corresponded to the 291 best alignments (left) and the <0.8 group corresponded to the 48 worst alignments. **(E-F)** Distributions describing the mean pLDDT and protein sequence length of AlphaFold structures that either (1) had analog MODELLER structures (red) or (2) did not (blue).

**Figure S5.**
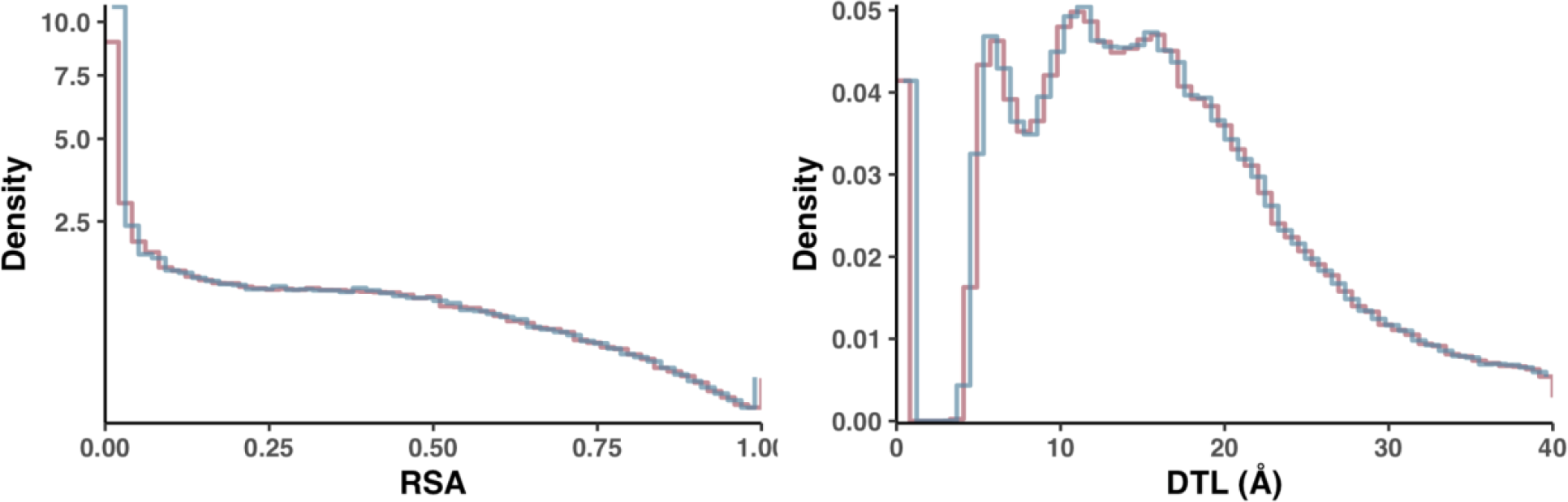
Comparison of null distributions for pN^(site)^ and pS^(site)^ for RSA and DTL. Each distribution was calculated by averaging 10 independent, randomly shuffled datasets of either pN^(site)^ (red line) or pS^(site)^ (blue line). To better visualize differences between the null distributions, the blue lines depicting the pS^(site)^ distributions were shifted right by half of a bin’s width.

**Figure S6.**
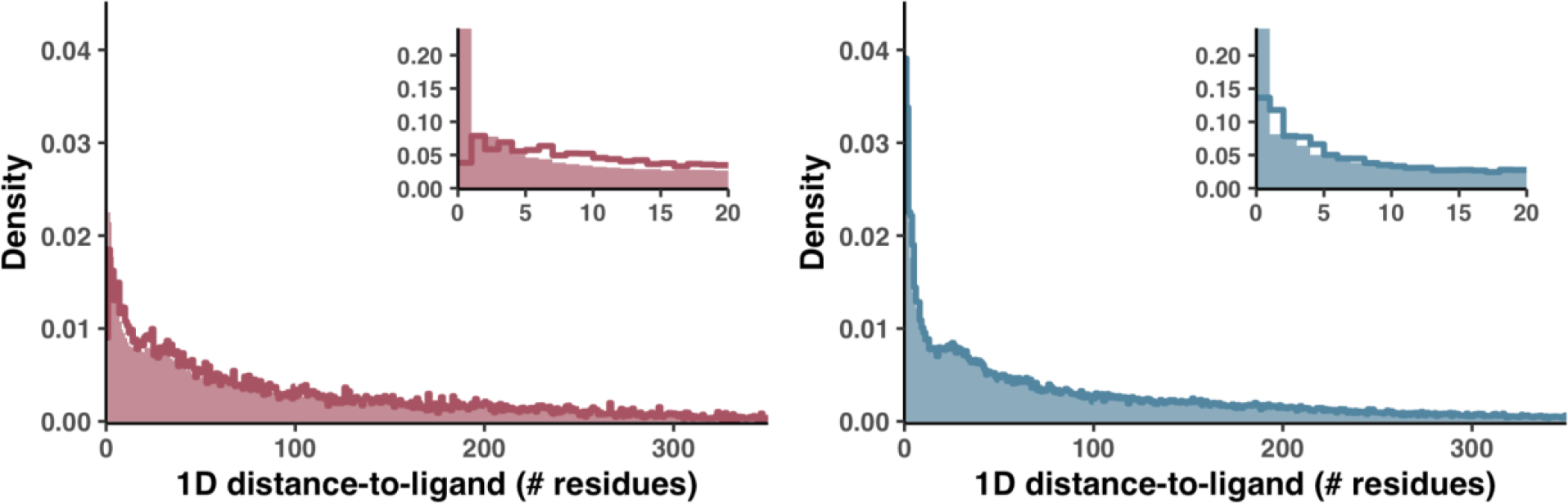
Functional constraint is less resolved when using a sequence-distance metric of DTL. pN^(site)^ (left panel) and pS^(site)^ (right panel) distributions with respect to 1D DTL, which we defined as the number of sites in a protein’s sequence that separate a given site from a predicted ligand-binding site. Lines represent the observed distributions, and filled regions represent the null distributions, calculated via the shuffling procedure described in Figure 2. Insets show the same data zoomed into the 1D DTL range [0, 20].

**Figure S7.**
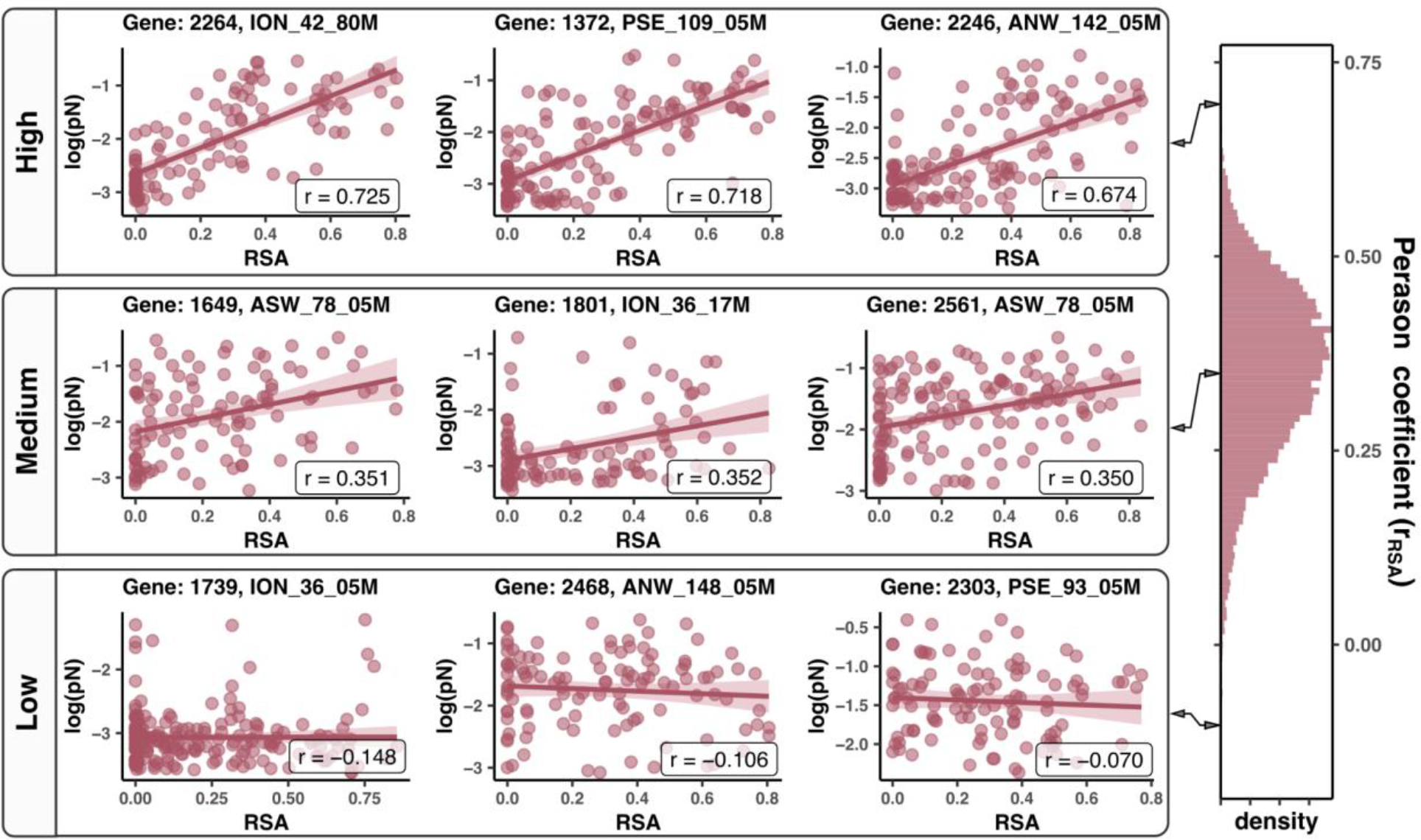
Select gene-sample pairs illustrate the diversity with which pN^(site)^ associates with RSA. Scatterplots for handpicked gene-sample pairs are shown from three regimes of model quality: high (top), mid (middle), and low (bottom). The right panel shows the distribution of Pearson coefficients, and the bin that each example was taken from is highlighted in pink. Each scatter plot is a gene-sample pair, each datapoint is a residue, the x-axis is the RSA of the residue, and the y-axis is the observed log10(pN^(site)^). Lines of best fit are shown in red, with 95% confidence intervals visualized translucently. The Pearson coefficients of each fit are labeled on the scatterplot.

**Figure S8.**
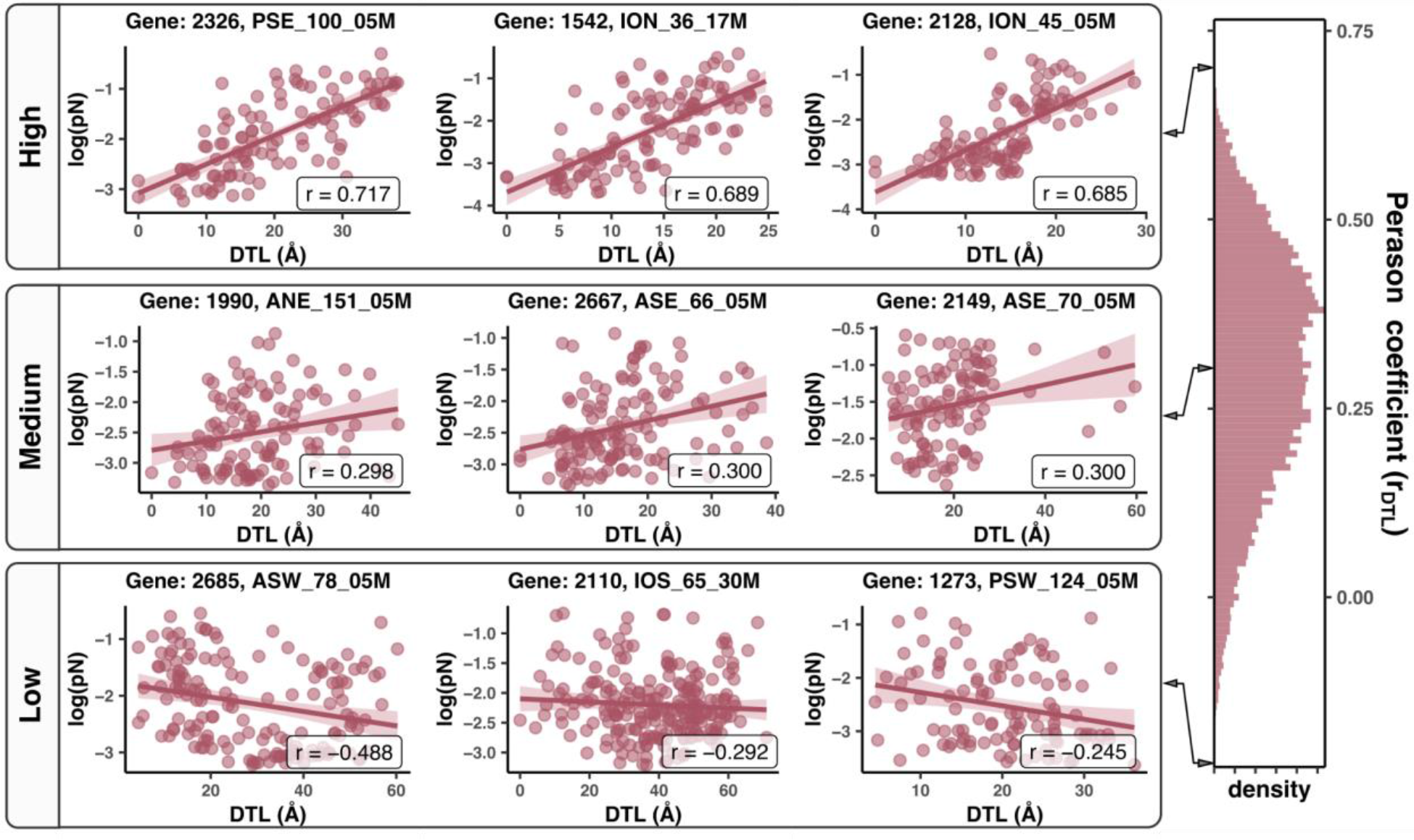
Select gene-sample pairs illustrate the diversity with which pN^(site)^ associates with DTL. Scatterplots for handpicked gene-sample pairs are shown from three regimes of model quality: high (top), mid (middle), and low (bottom). The right panel shows the distribution of Pearson coefficients, and the bin that each example was taken from is highlighted in pink. Each scatter plot is a gene-sample pair, each datapoint is a residue, the x-axis is the DTL of the residue, and the y-axis is the observed log10(pN^(site)^). Lines of best fit are shown in red, with 95% confidence intervals visualized translucently. The Pearson coefficients of each fit are labeled on the scatterplot.

**Figure S9.**
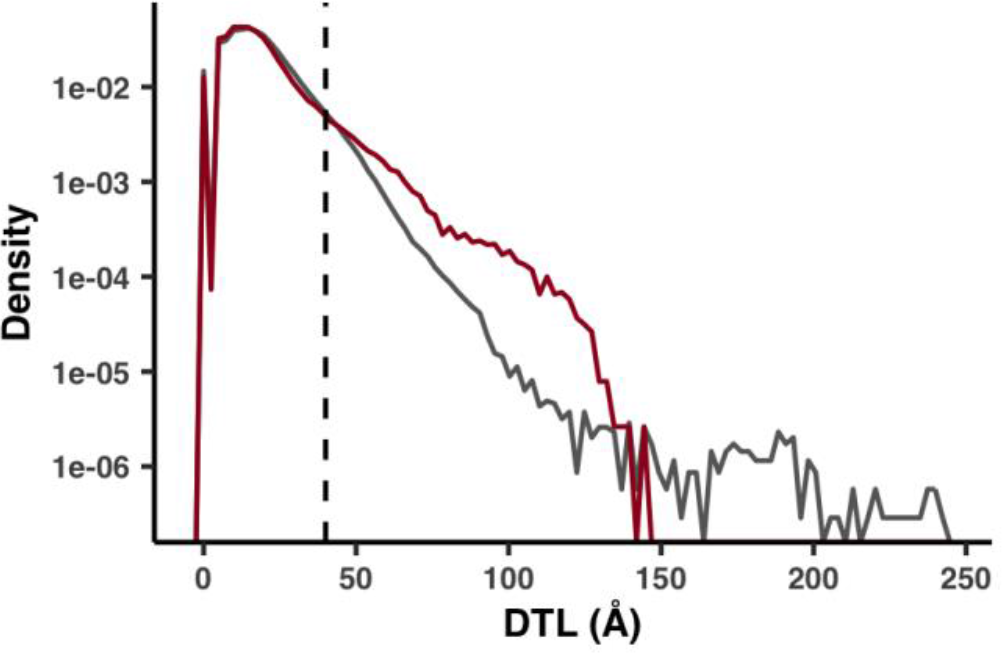
Incomplete ligand characterization leads to erroneously high DTL values. A comparison of DTL distributions (semi-log axis) for the 1a.3.V and the BioLiP database. The 1a.3.V core distribution (red) was calculated from all sites in the subset of genes with both a predicted structure and at least one predicted ligand-binding residue. The BioLiP distribution (gray) was calculated from the sites of 5,000 structures in the BioLiP database. For the 1a.3.V core, DTL was calculated as the distance to the closest predicted ligand-binding residue. For BioLiP, it was calculated as the distance to the closest annotated ligand-binding residue. For both methods, distance was calculated between the sites’ side chain center of masses. The dashed line marks the 40Å cutoff we used for all analyses besides Figure 2b, which excludes 8.0% of the total sites.

**Figure S10.**
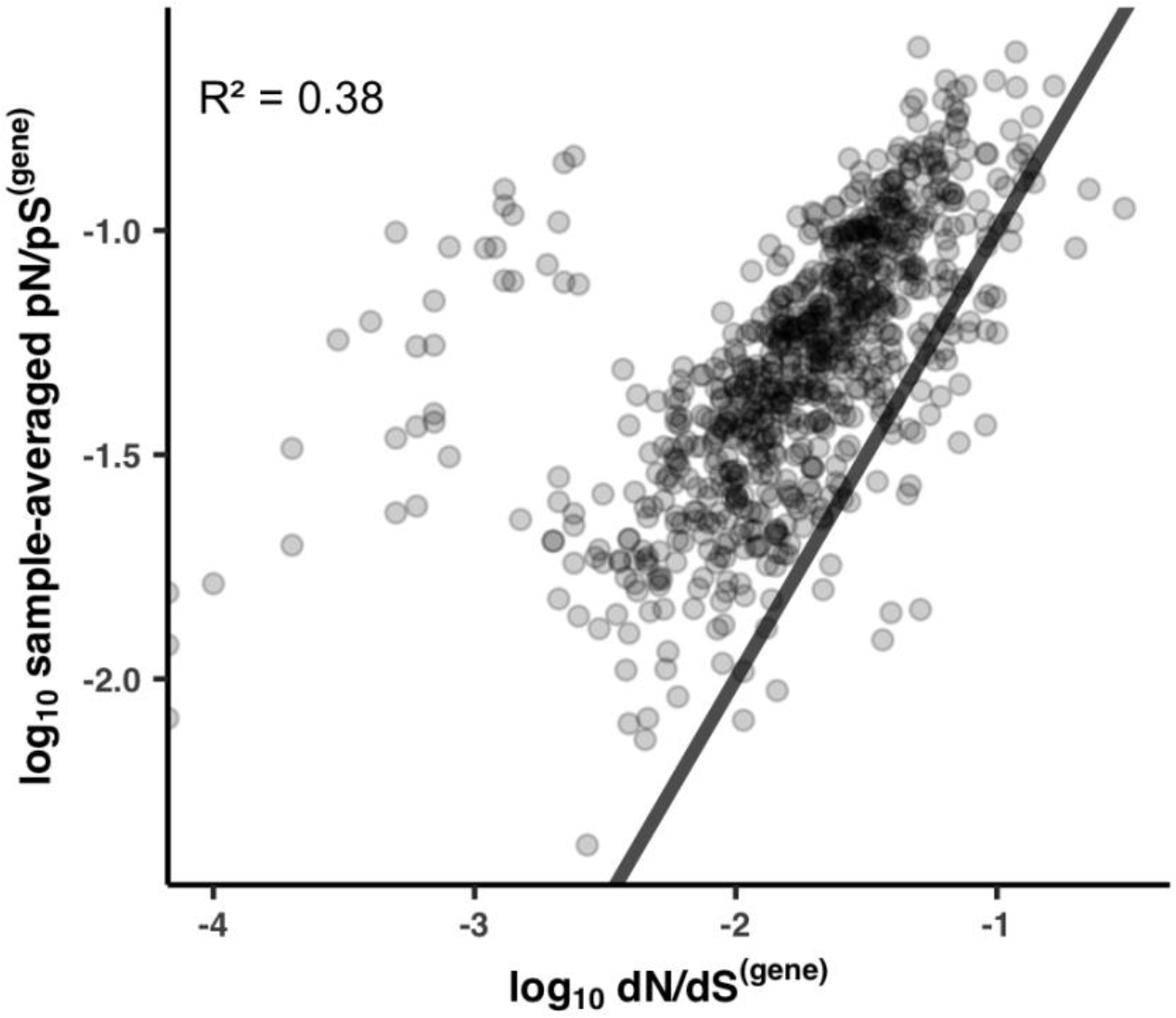
Sample-averaged pN/pS^(gene)^ values correlate with dN/dS^(gene)^ values between HIMB83 and HIMB122. The x- and y-axes are the log-transformed dN/dS^(gene)^ and sample-averaged pN/pS^(gene)^ values (respectively) for the 743 genes that (1) belonged to the 1a.3.V core and (2) had HIMB122 homologs. The black line is the equation y = x, meaning that genes above this line maintain sample-averaged pN/pS^(gene)^ values that exceed dN/dS^(gene)^. The R^2^ is for a linear regression of the log-transformed variables.

**Figure S11.**
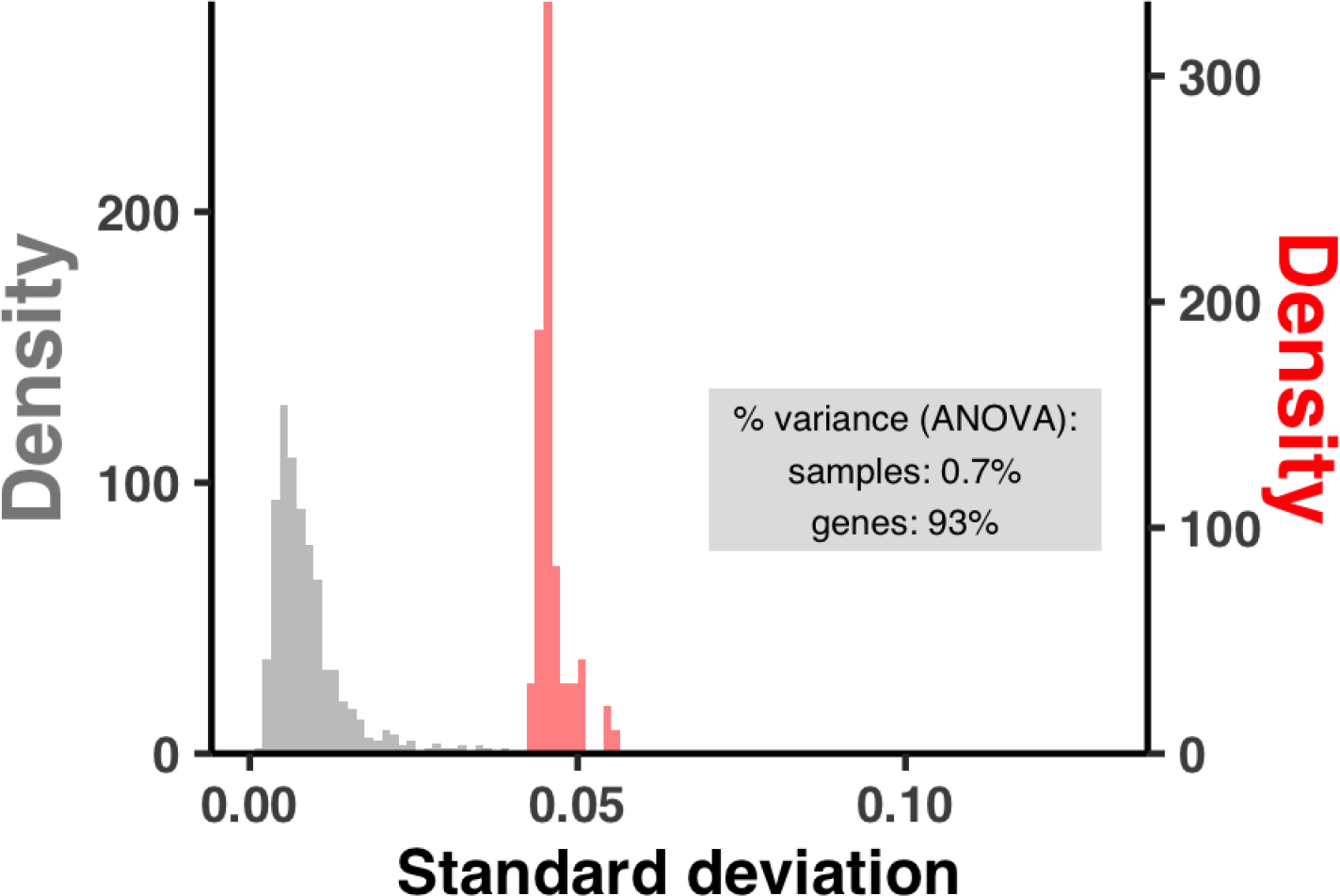
pN/pS^(gene)^ varies more significantly between genes in a given sample than between samples for a given gene. The x-axis is the standard deviation of either a sample’s pN/pS^(gene)^ values observed over genes (orange), or a gene’s pN/pS^(gene)^ values observed over the 74 samples (gray). The gray box denotes the amount of variance explained by genes and samples in an ANOVA from the linear model pN/pS^(gene)^ ∼ gene + sample.

**Figure S12.**
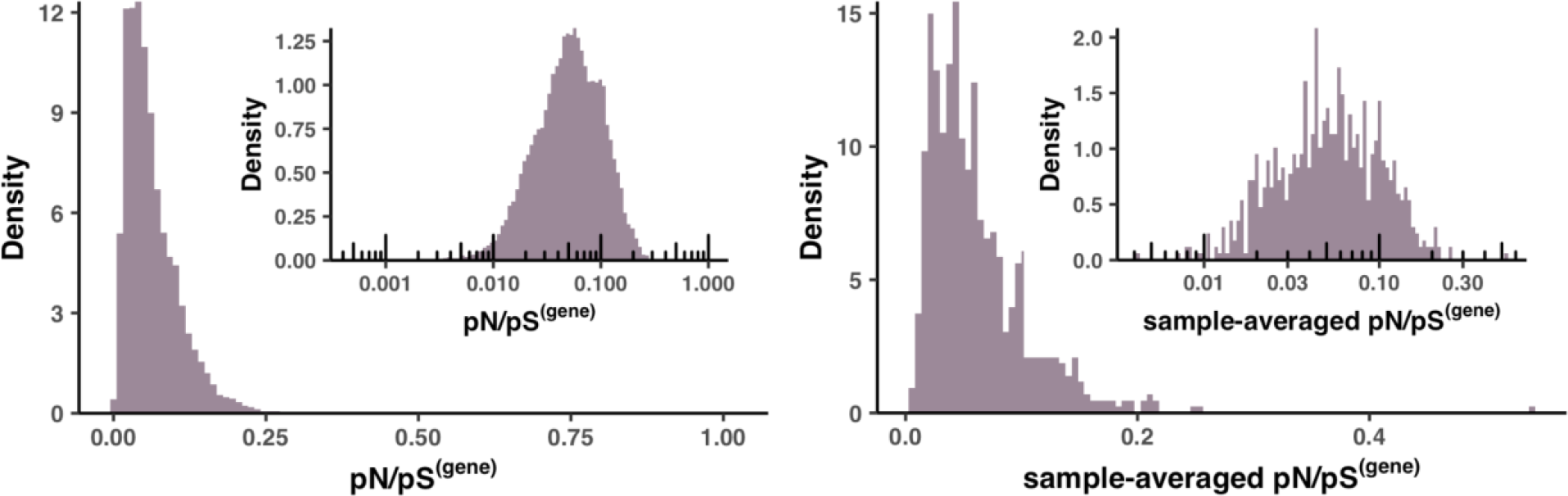
Distributions of pN/pS^(gene)^. Left panel shows the distribution of pN/pS^(gene)^, and the right panel shows the distribution of sample-averaged pN/pS^(gene)^. Insets show the same distributions with a log10-transformed x-axis.

**Figure S13.**
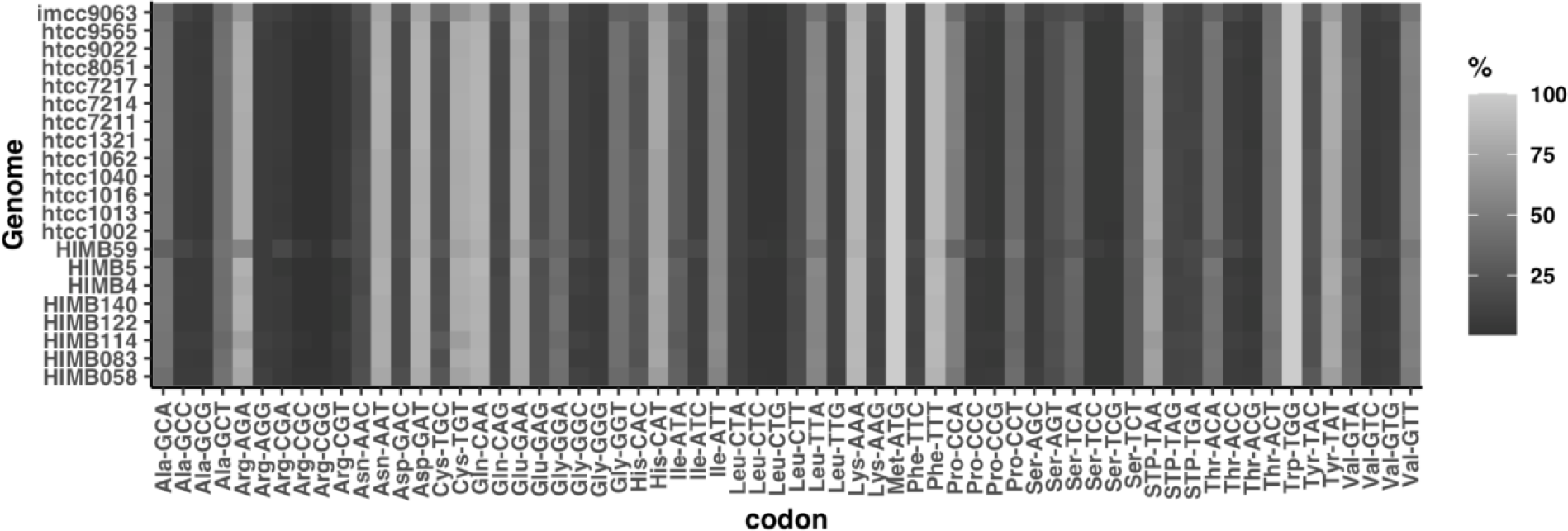
Codon usage of HIMB83 and 20 other genomes in the SAR11 clade.

**Figure S14.**
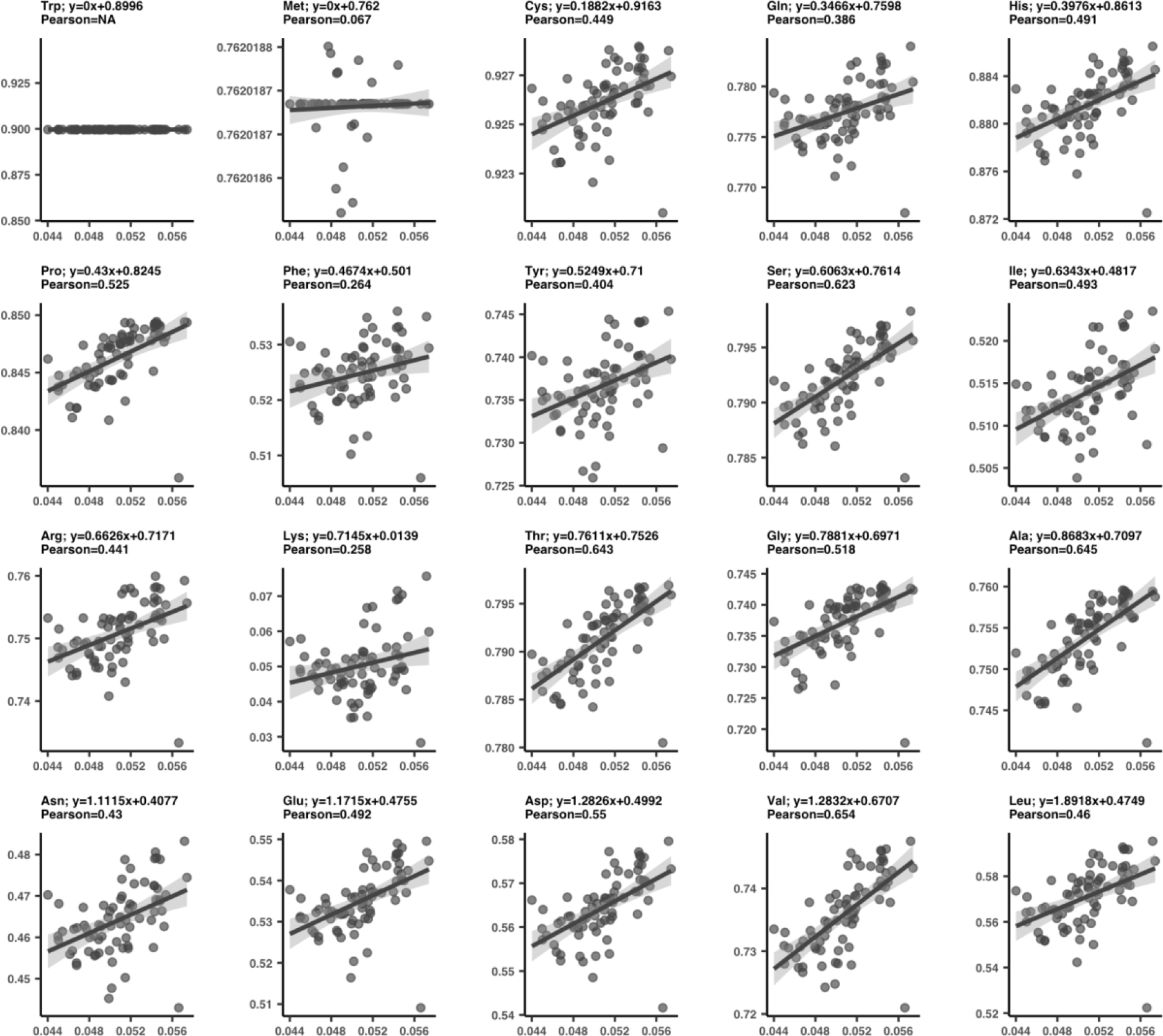
Codon rarity measured for each amino acid reveals varied response to selection strength, with most amino acids preferring rare codons in high selection samples. Each plot is a different amino acid, and each datapoint is a sample. The x-axis is pN/pS^(core)^, *i.e.* the ratio of nonsynonymous to s-polymorphism rates in the 1a.3.V core genome, and is shared between all plots. For a given plot, the y-axis was determined by first subsetting the polymorphism data to only include synonymous sites (in this instance we define synonymous as exhibiting pN^(site)^ < 0.0005) that corresponded to the given amino acid. Using lysine as an example, this led to on average 21,127 sites per sample. For each amino acid in each sample, we then calculated the overall codon rarity (y-axis) by averaging codon rarities across all included positions. A line of best fit (gray line) with 95% confidence intervals (light gray) is shown for each plot, with equation and Pearson correlation coefficient shown above.

