## Supplementary Information for "Structure-informed microbial population genetics elucidate selective pressures that shape protein evolution"

[1]Department of Medicine, University of Chicago, Chicago, IL, USA; [2] Graduate Program in Biophysical Sciences, University of Chicago, Chicago, IL, USA; [3]Department of Biostatistics, University of Washington, Seattle, WA, USA; [4]Hawai'i Institute of Marine Biology, University of Hawai'i at Mānoa, Kāne'ohe, Hawai'i, USA; [5]Department of Biochemistry & Molecular Biology, University of Chicago, Chicago, IL, USA; [6]Josephine Bay Paul Center for Comparative Molecular Biology and Evolution, Marine Biological Laboratory, Woods Hole, MA 02543, USA; [7]Helmholtz Institute for Functional Marine Biodiversity, Oldenburg, Germany; [8]Institute for Chemistry and Biology of the Marine Environment, University of Oldenburg, Oldenbug, Germany.

#### Regimes of sequence similarity probed by metagenomics, SAR11 cultured genomes, and protein families

We investigated how sequence similarity between HIMB83 and aligned metagenomic reads compares to the traditional methods of sequence comparisons between other SAR11 cultured genomes, as well as between members of associated protein families. To do this, we calculated the percent similarity (PS) between HIMB83 genes and (a) all aligned reads, (b) homologs found in 20 SAR11 ocean isolates, and (c) members of the best matching Pfam protein family.

For (a), PS values for each gene were calculated by considering one metagenome at a time. In each metagenome, the reads that aligned to the gene were captured, trimmed (so there were no reads overhanging the gene), and compared to the aligned segment of HIMB83. The PS was calculated by comparing non-gap positions. This was then averaged to yield a PS value for each gene-metagenome pair. To define a single PS value for each gene, PS values were averaged across metagenomes.

For (b), gene clusters were calculated for HIMB83 and 20 additional SAR11 isolates using the anvi'o pangenomic workflow. An MSA was built from the sequences of each gene cluster using muscle (Edgar 2004), and then each non-HIMB83 sequence was compared to the HIMB83 sequence. The PS was determined by calculating the fraction of matches in non-gap positions. Each HIMB83 gene was attributed a single PS value by averaging PS values in each pairwise comparison, weighted by the number of non-gap positions in the pairwise alignment. Gene clusters containing multiple HIMB83 genes were ignored.

For (c), HIMB83 genes were matched to Pfam protein families via the anvi'o program `anvi-run-pfams`. Hits that passed the GA gathering threshold were retained, and the best hit (lowest e-value) for each HIMB83 gene was defined as the associated Pfam. For each HIMB83 gene, the associated Pfam seed sequence MSA was downloaded using the Python package prody (S. Zhang et al. 2021) and the HIMB83 protein sequence was added to the MSA using muscle. PS values were calculated from the MSAs in a manner identical to that outlined in (b). It is important to note that this comparison used protein sequences, whereas (a) and (b) both used nucleotide sequences.

Figure S1 shows the distribution of percent similarities for each comparative method, roughly indicating the distinct regimes of evolutionary relatedness that each method probes. Unsurprisingly, protein families are most evolutionarily divergent (mean amino acid PS 28.8%). Relative to SAR11 homologs (mean nucleotide PS 77.3%), the aligned reads are highly related (mean nucleotide PS 94.5%), showing that metagenomics offers a modality of sequence inquiry more highly resolved than sequence comparisons between isolated cultures.

### Comparing structure predictions between AlphaFold and MODELLER

The biggest difference between structure prediction methods was the expectedly higher portion of predictions yielded by AlphaFold. While AlphaFold produced 754 structures we deemed trustworthy (see Methods), MODELLER produced 346 due to its reliance on pre-existing template structures. In 339 cases both methods procured a structure prediction for a given protein sequence, and it is within this intersection that we drew comparisons between the methods' structures.

We compared the topological similarity between AlphaFold and MODELLER structures using TM score (Y. Zhang and Skolnick 2004) and alpha carbon RMSD. Overall, the distributions of these metrics (Figures S4a, S4b) illustrate the overarching similarity between AlphaFold and MODELLER structures. Poor RMSD scores were usually the result of multidomain proteins linked by unstructured chains, and not the result of large structural discrepancies. TM scores better handle these cases. Since a TM score of 0.5 indicates that proteins likely belong to the same fold family (Xu and Zhang 2010), our average TM score of 0.88 indicates strong overall agreement between AlphaFold and MODELLER.

On average, AlphaFold yielded a higher proportion of secondary structure (Figure S4c), and we found this discrepancy to be most pronounced when TM scores were low (<0.8) (Figure S4d). In fact, for the worst alignments (TM score <0.6), in 15 of 16 cases AlphaFold yielded more secondary structure.

Next, we turned our attention to proteins that AlphaFold predicted structures for, but that MODELLER did not due to absent templates. These proteins were on average smaller (Figure

S4e) and yielded lower mean pLDDT scores compared to structures possessing a MODELLER analog. Since AlphaFold is trained on pre-existing structures, this result is expected and lends credence to pLDDT as a metric for fold confidence. Even still, these structures averaged a mean pLDDT score of 90.8, which is considered to be highly accurate (Jumper et al. 2021).

Overall, our findings suggest that overall similarity between the two methods is high, that AlphaFold may be outperforming MODELLER due to increased fraction of secondary structure, and that proteins modeled by AlphaFold but not MODELLER are still considered highly accurate predictions.

#### RSA and DTL predict nonsynonymous polymorphism rates

To complement our analyses in which we estimated the percentage of polymorphism data that can be explained by RSA and DTL (Table S6, Methods), we constructed synonymous models (s-models) and nonsynonymous model (ns-models) for each gene in each sample. We excluded monomorphic sites ($pN^{(site)} = 0$ for ns-models, $pS^{(site)} = 0$ for s-models), sites with DTL > 40Å (see Methods), and removed gene-sample pairs containing <100 remaining sites, resulting in 16,285 ns-models and 24,553 s-models (Table S7).

We fit linear models of $\log_{10}(pN^{(site)})$ and $\log_{10}(pS^{(site)})$ to RSA. We found that applying a logarithmic function to polymorphism rates yielded better fits than without. We filtered out any genes that did not have a predicted structure and at least one predicted ligand-binding site, which when applied in conjunction with the above filters resulted in 381 genes for the s-models and 342 genes for the ns-models. ns-models yielded consistently positive correlations (average Pearson coefficient of $r_{RSA} = 0.353$) (Figure 2c), whereas s-models exhibited correlations centered around 0 (average $r_{RSA} = -0.029$). The average $R^2$ was 0.137 for ns-models, however model quality varied significantly between gene-sample pairs. In fact, we found that $R^2$ varied from as high as 0.526 (gene 2264 in sample ION_42_80M), to as low as 0.0% (gene 2486 in sample ION_42_80M). Lines of best fit for select gene-sample pairs illustrate the range of correlatedness seen between $\log_{10}(pN^{(site)})$ and RSA (Figure S7). Overall, these results show that RSA is a significant predictor that partially explains the differences in polymorphism rates observed between sites in a given gene and sample.

Using the same procedure, we linearly regressed $\log_{10}(pN^{(site)})$ and $\log_{10}(pS^{(site)})$ with DTL and found that 96% of ns-models yielded positive correlations with DTL with considerable predictive power, where on average 11.5% of per-site ns-polymorphism rate variation could be explained by DTL (Table S7). $R^2$ values varied significantly, ranging from 0.514 (gene 2326 in sample PSE_100_05M) to 0.0% (gene 2246 in sample PSE_102_05M). Lines of best fit for select gene-sample pairs illustrate the range of relatedness observed between $\log_{10}(pN^{(site)})$ and DTL (Figure S8). Interestingly, we found that $\log_{10}(pS^{(site)})$ on average negatively correlates with DTL (average Pearson coefficient -0.057). The overall positive correlation of DTL with $\log_{10}(pN^{(site)})$ suggests that on a proteomic scale, selection for function imposes a spectrum of per-site selective pressures, where pressure increases with proximity to ligand-binding regions.

Individually, RSA and DTL respectively explain 13.7% and 11.5% of per-site ns-polymorphism rate variance. To quantify their collective explanatory power, we fit a third set of models that linearly regressed $\log_{10}(pN^{(site)})$ and $\log_{10}(pS^{(site)})$ with RSA and DTL together (Figure SI2; Table S7). A Pearson correlation between RSA and DTL revealed the relative independence of each variable from the other ($R^2$ = 0.082, r = 0.286), precluding effects of multicollinearity (Figure SI1). The results revealed that including both RSA and DTL yielded a considerably better set of models for ns-polymorphism rates, with an average explained variance of 17.7% (average adjusted $R^2_{RSA\text{-}DTL} = 0.177$).

The predictive power of RSA and DTL illuminates how structural and functional constraints influence polymorphism rates by shaping the confines within which neutral evolution operates (Worth, Gong, and Blundell 2009), yet observed rates can also be dominantly driven by stochastic processes of mutagenesis and drift. For example, no site will be polymorphic in the absence of a seeding mutagenesis event, even if under low structural and functional constraints. Thus, polymorphism rates are determined in part by constraints, and in part by random chance, the latter of which diminishes the predictive power of RSA and DTL when modeling polymorphism rates of individual sites.

By averaging across groups of sites, we vastly increased the signal-to-noise ratio of polymorphism rate data and revealed a two parameter model (RSA and DTL) that explains the majority of ns-polymorphism trends. To reduce per-site noise, we first grouped sites sharing similar RSA and DTL values so that each group contained the same order of magnitude of data (axes in Figure 2e, Table S8). For example, the group ($RSA_1$, $DTL_2$) contains the 3,164 sites with RSA values in the 1st RSA range [0.00,0.01) and DTL values in the 2nd DTL range [5.0Å,6.4Å). Then, we calculated per-group polymorphism rates $pN^{(group)}$ and $pS^{(group)}$, which are weighted averages of $pN^{(site)}$ and $pS^{(site)}$ values found within a group (see Methods). Averaging polymorphism rates across sites that exhibit similar RSA and DTL values has the effect of averaging out per-site and per-sample variance, which we found to reveal impressive proteome-wide trends in polymorphism rates with respect to RSA and DTL. $pN^{(group)}$ values from each group collectively describe a 2D surface (Figure 2e, Table S8), where one axis illustrates how structurally constrained sites tend to be due to RSA and the other axis illustrates how functionally constrained sites tend to be due to DTL. In contrast to the noisy $pN^{(site)}$ data observed within gene-sample pairs (Figures S7, S8), the $pN^{(group)}$ surface is smooth and roughly linear (Figure 2e). Nonsynonymous polymorphism rates of groups varied from as low as 0.001 to as high as 0.021. A group's polymorphism rate appeared to be chiefly determined by the overall constraint of its sites, which is a composite of both structural and functional constraints. Structural and functional constraints appeared to be additive, such that sites with both low RSA and DTL (left panel of Figure 2e, bottom-left) statistically exhibited the lowest rates of ns-polymorphism, and sites with both high RSA and DTL (left panel of Figure 2e, top-right) statistically exhibited the highest rates of polymorphism. Additionally, these constraints are seen to act independently of one another: some groups exhibit low $pN^{(group)}$ due to structural constraint (top-left) while others exhibit low $pN^{(group)}$ due to functional constraint (bottom-right), illustrating that selection for structure and selection for function can independently constrain evolution.

Sites exhibited a spectrum of ns-polymorphism rates that is roughly linear. We determined this by fitting a linear model $pN^{(group)} \sim i + j$, where i refers to the group's RSA and DTL indices ($RSA_i$, $DTL_j$), yielded an adjusted $R^2$ of 0.836, meaning that 83.6% of ns-polymorphism rate variation can be explained by RSA and DTL when averaging over per-site effects (Figure SI3, Table S8). Increasing the number of groups decreased the number of sites in each group, weakening the efficacy of signal averaging, which expectedly decreased model quality. Even still, $R^2$ values for nonsynonymous models were robust to group numbers ranging from 4 (2x2) to 1,444 (38x38) (Figure SI4).

Site averaging yielded an unexpected relationship between s-polymorphism rates and RSA/DTL. $pS^{(group)}$ is not as strongly affected by RSA or DTL as $pN^{(group)}$, as indicated by the noisy contour lines of its surface (right panel Figure 2e). Even still, the linear model $pS^{(group)} \sim i +$

j yielded a significant, anti-correlated relationship with both RSA and DTL (adjusted $R^2$ of 0.206), in which s-polymorphism rates tended to decrease when RSA and DTL were high (Figure SI3). We have observed this surprising finding through other means as well: in the sample-gene models, (1) the mean Pearson correlation coefficient between $pS^{(site)}$ and RSA is -0.013 (Figure 2c), and (2) the mean Pearson correlation coefficient between $pS^{(site)}$ and DTL is -0.052 (Figure 2d). Signal averaging has revealed the extent of its effect: 20.6% of s-polymorphism rates can be explained by RSA and DTL when averaging over per-site effects, compared to 83.6% for ns-polymorphism rates.

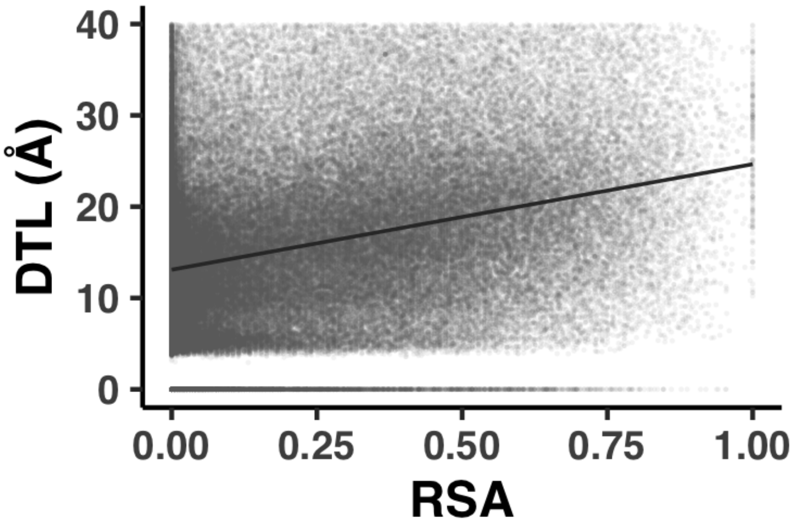


**Figure SI1. RSA and DTL are not problematically correlated.** Scatter plot of RSA vs. DTL for the 143,181 sites belonging to genes with a predicted structure and at least one predicted ligand. The line of best fit is shown in black, The Pearson coefficient is 0.313 and the $R^2$ is 0.098.

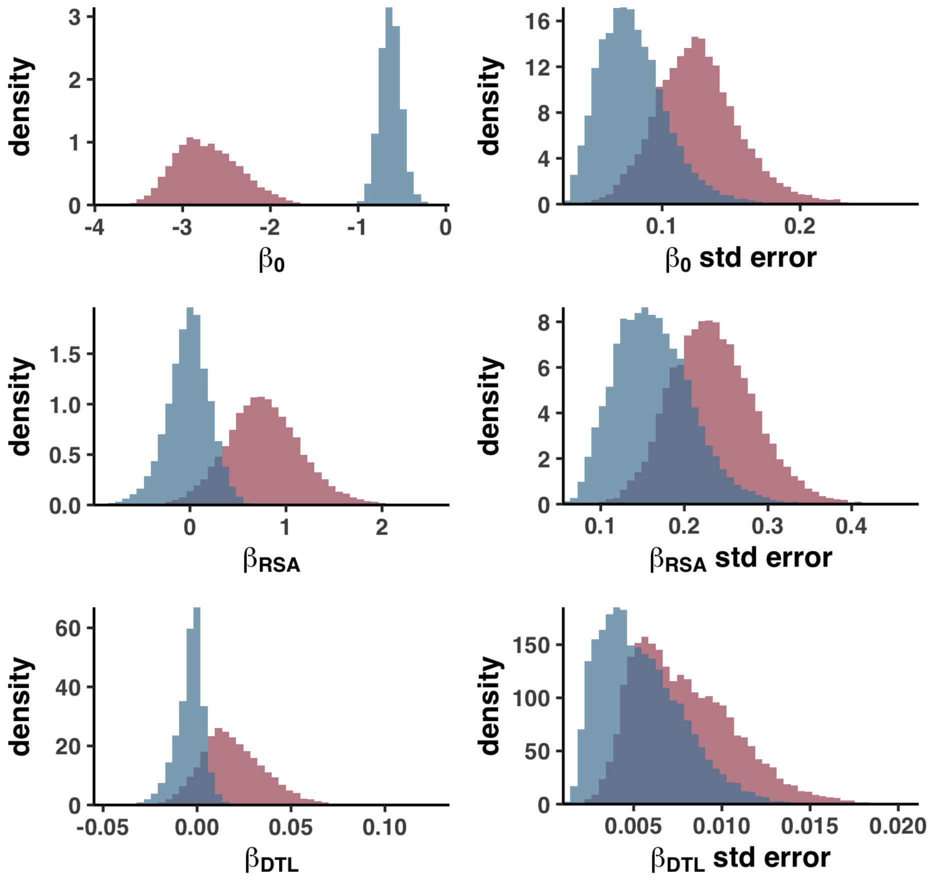


**Figure SI2. Parameter estimate and standard error distributions of the multidimensional linear regression models for $pN^{(site)}$ and $pS^{(site)}$.** Red denotes parameter/error distributions for the 16,285 nonsynonymous models of the form $pN^{(site)} = \beta_0 + \beta_{RSA}RSA + \beta_{DTL}DTL$ and blue denotes parameter/error distributions for the 24,553 models of the form $pS^{(site)} = \beta_0 + \beta_{RSA}RSA + \beta_{DTL}DTL$.

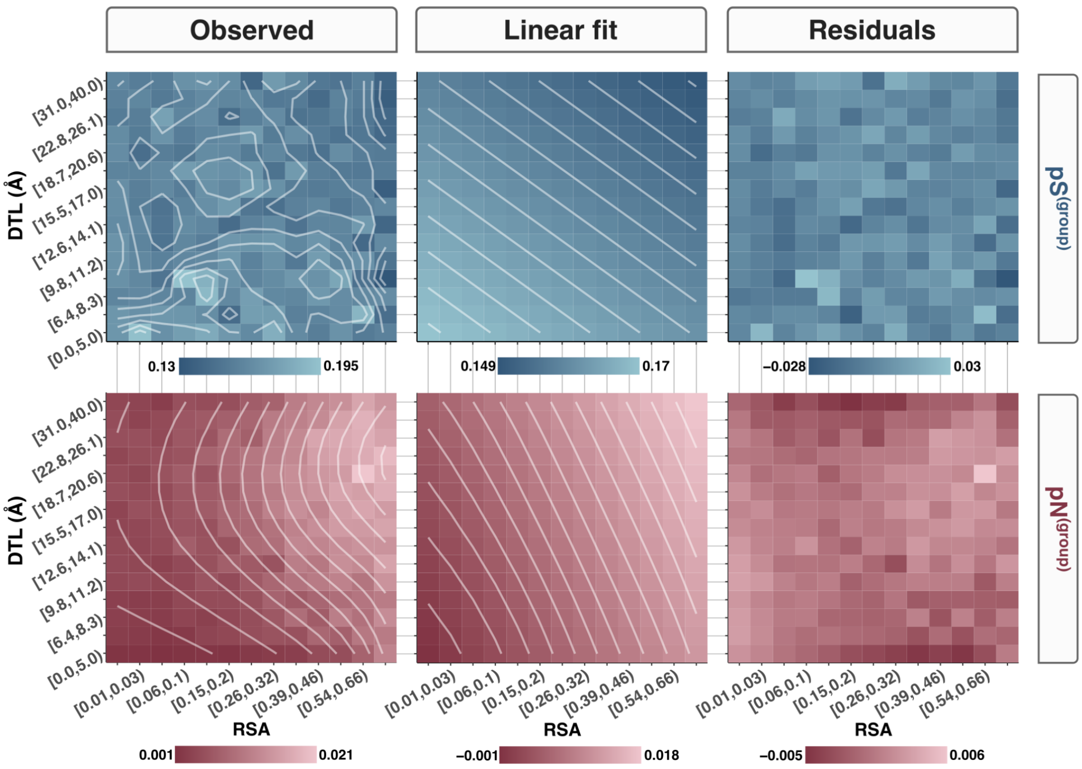


**Figure SI3. Observations, fits, and residuals of linear regressions for $pN^{(group)}$, $pS^{(group)}$, and $pN/pS^{(group)}$.** The x-axis and y-axis for each heatmap are RSA and DTL groups, respectively. The first column shows the observed values (those seen in Figure 2e), the second column shows the planes of best fit, and the third column shows the residuals. A legend for corresponding colors to values are shown below each heatmap. Contour lines for observed values and planes of best fit are shown as white and are calculated from smoothed data. Note that for the planes of best fit, the contour lines of the underlying data are by definition straight and perpendicular to one another, though due to edge effects of the smoothing procedure, there is a slight bend in the visualization of some contour lines.

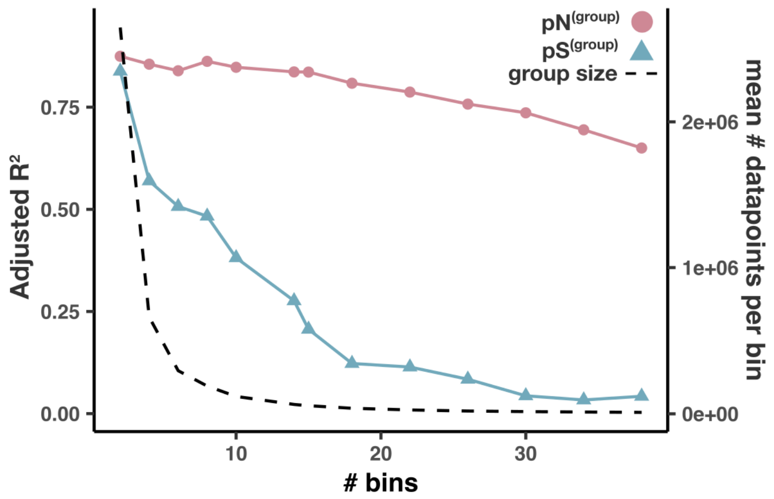


**Figure SI4. Model quality decreases for pN$^{(site)}$ and pS$^{(group)}$ as the number of RSA and DTL groups increases.** The x-axis represents how many bins RSA and DTL are each split into. For example, the heatmaps in Figure 2e correspond to # bins = 15, since RSA and DTL are split into 15 bins, totaling 225 (=15x15) groups. The left y-axis corresponds to the adjusted $R^2$ value for the models pN$^{(group)}$ (red) and pS$^{(group)}$ (blue). The right y-axis corresponds to the average number of data points (# sites multiplied by # samples) found in a group (dashed black line).

### dN/dS$^{(gene)}$ and sample-averaged pN/pS$^{(gene)}$ yield consistent results

To validate our pN/pS$^{(gene)}$ calculations, we ascribed a sample-averaged pN/pS$^{(gene)}$ value to each gene and compared the values to dN/dS$^{(gene)}$ (Table S12), a more commonly and classically utilized metric that is the ratio of nonsynonymous to synonymous substitutions observed between homologous genes of two or more species. We calculated dN/dS$^{(gene)}$ for 753 homologous gene pairs found between HIMB83 and a closely related cultured representative HIMB122 (see Methods). Importantly, ANI between HIMB83 and HIMB122 was 82.6%, whereas the average ANI between HIMB83 and recruited reads was 94.5%, making it unlikely that sample-averaged pN/pS$^{(gene)}$ and dN/dS$^{(gene)}$ were cross-contaminated due to HIMB83 recruiting significant proportions of reads from HIMB122-like populations. We found that log-transformed sample-averaged pN/pS$^{(gene)}$ highly correlated with log-transformed dN/dS$^{(gene)}$ (Pearson $R^2$ = 0.380), showing that the two metrics are commensurable. Nevertheless, differences were expected and observed. The ratio between sample-averaged pN/pS$^{(gene)}$ and dN/dS$^{(gene)}$ was on average 6.23 (Figure S10), matching expectations that slightly deleterious, nonsynonymous mutants commonly drift to observable frequencies, yet far less commonly drift to fixation.

#### Transcript abundance largely explains genic differences in the strengths of purifying selection

Sample-averaged pN/pS$^{(gene)}$ values varied significantly between genes, varying from 0.004-0.539, with a mean of 0.063 (Figure S12, Table S9). What causes such variation in purifying selection strengths? Across diverse taxa (Drummond and Wilke 2008), it has been shown that highly expressed proteins evolve more slowly due to being selectively constrained to be robust to mistranslation in order to safeguard against toxicity of misfolded proteins, whose detrimental fitness costs scale with expression level (Drummond et al. 2005). We assessed the extent to which expression level may explain purifying selection variation in 1a.3.V by calculating metatranscriptomic coverage values for each 1a.3.V core gene in the 50 of 74 environments that had accompanying metatranscriptomics datasets (see Methods). We defined transcript abundance (TA) as the ratio of metatranscriptomic to metagenomic relative abundances (see Methods), which yielded a widely skewed distribution of values (Figure SI5a, Table S13).

Comparing sample-median TA values to sample-averaged pN/pS$^{(gene)}$ values yielded a strong, negative correlation (Figure SI5b, Pearson r = -0.539, $R^2$ = 0.290) according to an inverse power-law relationship. The specific form of the linear model used was $\log_{10}(\text{median}_s(\text{TA})+0.01) \sim \log_{10}(\text{mean}_s(\text{pN/pS}^{(gene)}))$, where $\text{median}_s$ and $\text{mean}_s$ denote the median and mean across samples for a given gene, respectively. To avoid excluding zeros, we added 0.01 to the log-transformation of $\text{median}_s$(TA). These findings indicate that 29.0% of purifying selection variation between genes can be explained via transcript abundance alone, a value in line with what has been observed between yeast homologs (Drummond et al. 2005). Overall, these results recapitulate a central result in protein evolution, and demonstrate its validity *in situ* using culture-independent approaches that link genetic variation and transcript abundance for a naturally occurring microbe.

Next, we tested whether $pN/pS^{(gene)}$ values between samples of a given gene also follow an inverse power-law relationship with TA. We found that of the 799 genes tested, 74% exhibited (weak) negative correlations between $\log_{10}$(TA+0.01) and $\log_{10}(pN/pS^{(gene)})$ (Figure SI5c), yet only 11.5% of genes passed significance tests (one-sided Pearson, 25% Benjamini-Hochberg false discovery rate) (Figure SI5d). Given the strong correlation observed between genes, the lack of correlation observed between samples is a seemingly contradictory result, yet can be attributed to a difference in timescales: TA fluctuates on the order of minutes, often occurring in 'bursts', whereas $pN/pS^{(gene)}$ is shaped over time scales orders of magnitude longer than the 2 week replication time of SAR11. Since metagenome-metatranscriptome pairs sample single snapshots in time, measured TAs are unlikely to reflect the time-averaged values that constrain $pN/pS^{(gene)}$. These fluctuations therefore muddy signals that may exist between $pN/pS^{(gene)}$ and TA. Smoothing these fluctuations by averaging across samples thereby reveals the strong negative correlation observed (Figure SI5b). In other words, TAs that are not averaged across environments are unreliable proxies for overall transcription level.

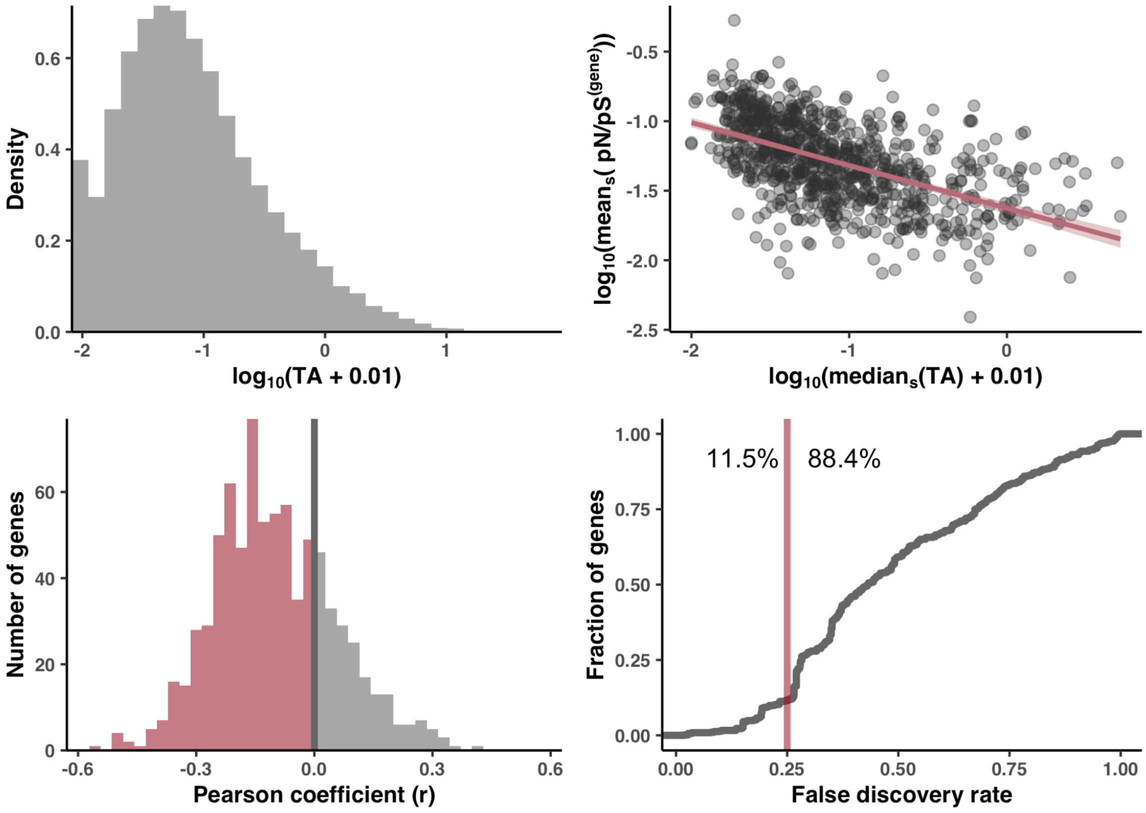


**Figure SI5. Associations of transcript abundance (TA) data with pN/pS$^{(gene)}$. (A) Log-transformed distribution of TA values across genes and samples.** See Methods for details on TA calculation. 0.01 has been added to the log-transformation to avoid the exclusion of zeros. **(B) TA is a strong predictor of pN/pS$^{(gene)}$ when pooling data across samples.** Each datapoint is a gene, where the x-axis is the gene's median TA across samples, the y-axis is the gene's sample-averaged pN/pS$^{(gene)}$, and each axis has been log-transformed. The linear model yielded a Pearson coefficient of 0.539, an $R^2$ of 0.290 and a line of best fit y = (-0.31 ± 0.02)x + (-1.63 ± 0.02) shown in pink (95% confidence intervals shown in translucent pink). **(C) pN/pS$^{(gene)}$ between samples of a given gene weakly correlate (on average) with TA.** A one-side Pearson correlation between $\log_{10}$(TA + 0.01) and $\log_{10}$(pN/pS$^{(gene)}$) was calculated separately for 799 genes, resulting in the following distribution of Pearson coefficients, of which 74% were negative (pink). **(D) Accounting for multiple testing yields few statistically significant negative correlations.** The x-axis is the Benjamini-Hochberg false discovery rate (FDR) and the y-axis is the fraction of genes that have statistically meaningful negative correlations for a given FDR. Allowing a FDR of 25% (pink line), only 11.5% of genes have statistically significant negative correlations of $\log_{10}$(TA + 0.01) with $\log_{10}$(pN/pS$^{(gene)}$).

#### Stability analysis of polymorphism distributions with respect to pN/pS$^{(core)}$

To assess whether the ‘use it or lose it’ accumulation of ns-polymorphism in low RSA/DTL sites was specific to GS, or a more general feature of 1a.3.V, we performed a comparable procedure where instead of restricting our analysis to GS, we compiled polymorphism rates across all sites in genes with predicted structures and ligand-binding sites, and calculated pN/pS$^{(core)}$ for each sample, which serves as a proxy for genome-wide selection strength (see Methods). Within this dataset, we observed the same phenomena: in samples with high selection strength (high pN/pS$^{(core)}$), ns-polymorphism throughout the genome distributed (a) in more solvent-exposed sites (Figure 4a) and (b) farther from predicted binding sites (Figure 4b). Our bootstrapping stability analysis (Figure SI6, Table S14) showed that in 99.5% of gene resamplings, the mean RSA of ns-polymorphism negatively associated with pN/pS$^{(core)}$ (one-sided Pearson coefficient p-value <0.05), whereas in only 69.5% of gene resamplings did the mean DTL of ns-polymorphism negatively associate with pN/pS$^{(core)}$. This latter finding indicates that the signal in Figure 4b is driven by an incomplete set of the 1a.3.V core genes. We hypothesized this is due to the many shortcomings of DTL estimation discussed priorly leading to false-positive and/or false-negative ligand predictions that skew DTL distributions, or that not all ligands constraint ns-polymorphism patterns equally.

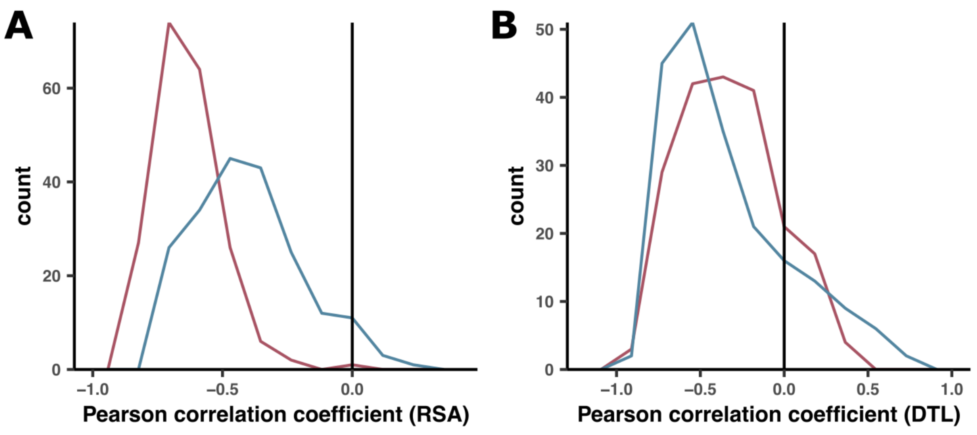


**Figure SI6. Robustness of negative associations between sample selection strength (pN/pS$^{(core)}$) and mean RSA/DTL of polymorphisms.** We tested the robustness of results in Figure 4 by performing a bootstrapping stability analysis in which we created 200 bootstrapped estimates of the correlation coefficients, where each bootstrap was a resampling of genes. **(A)** Histograms of the correlation coefficients between the mean RSA of s-polymorphism (blue) and ns-polymorphism (red) versus pN/pS$^{(core)}$. These correspond to Figures 4a and 4c, respectively. **(B)** Histograms of the correlation coefficients between the mean DTL of s-polymorphism (blue) and ns-polymorphism (red) versus pN/pS$^{(core)}$. These correspond to Figures 4b and 4d, respectively.

### Enabling interactive, exploratory, structure-informed metagenomic analyses using anvi-display-structure

There is an absence of computational tools that allow researchers to interactively explore metagenomic sequence variance in the context of predicted protein structures and ligand-binding sites. We addressed this gap by developing an interactive interface in which users can visualize, filter, and interact with metagenomic sequence variants in the context of modeled protein structures and predicted binding sites (Figures SI7, SI8, SI9, SI10). The exploratory analyses enabled by the interface is what has made the current research possible.

We created an interactive interface that dynamically processes data from anvi’o databases, which is done with the program `anvi-display-structure`. Once the interactive interface is initiated, users can select any gene with a modeled structure in their dataset, upon which anvi'o renders the predicted structure of the gene using NGL (Rose and Hildebrand 2015; Rose et al. 2016) and overlays sequence variants from metagenomes directly on the structure. By default, all variants across all metagenomes for a given gene are superimposed on a single display, however, the user can subdivide the display into as many as 16 sub-displays to compare and contrast variation across arbitrary groups of metagenomes (Figure SI7). The interface offers numerous ways to interact with and explore single-codon variants (SCVs) and single-amino acid variants (SAAVs). Hovering the mouse above any variant reveals its allele frequency vector and structural information of the reference residue such as solvent accessibility and secondary structure (Figure SI8). Interactive sliders filter variants displayed on structures through a suite of continuous, discrete, and categorical variables, including variant-specific parameters such as site entropy, solvent accessibility, BLOSUM scores of the competing alleles, residue number, and secondary structure (Figure SI8). These same variables can also dynamically change the color and size of individual variants (Figure SI9). Filters can be combined for exploratory investigations. For example, a user could simultaneously color variants by site entropy, size them by their coverage in metagenomes, and filter out those that exhibit high solvent accessibility (Figure SI9). The protein surface and backbone can be colored according to arbitrary user-provided data, for example, to visualize predicted binding sites of the protein. `anvi-display-structure` can save and load sessions to preserve filters, export displays as PNG images, and generate rich tabular outputs for allele frequencies and other properties of displayed variants. Finally, users can faithfully migrate the current view into PyMOL (Schrödinger, LLC) for further graphical refinement or statistical analyses (Figure SI10).

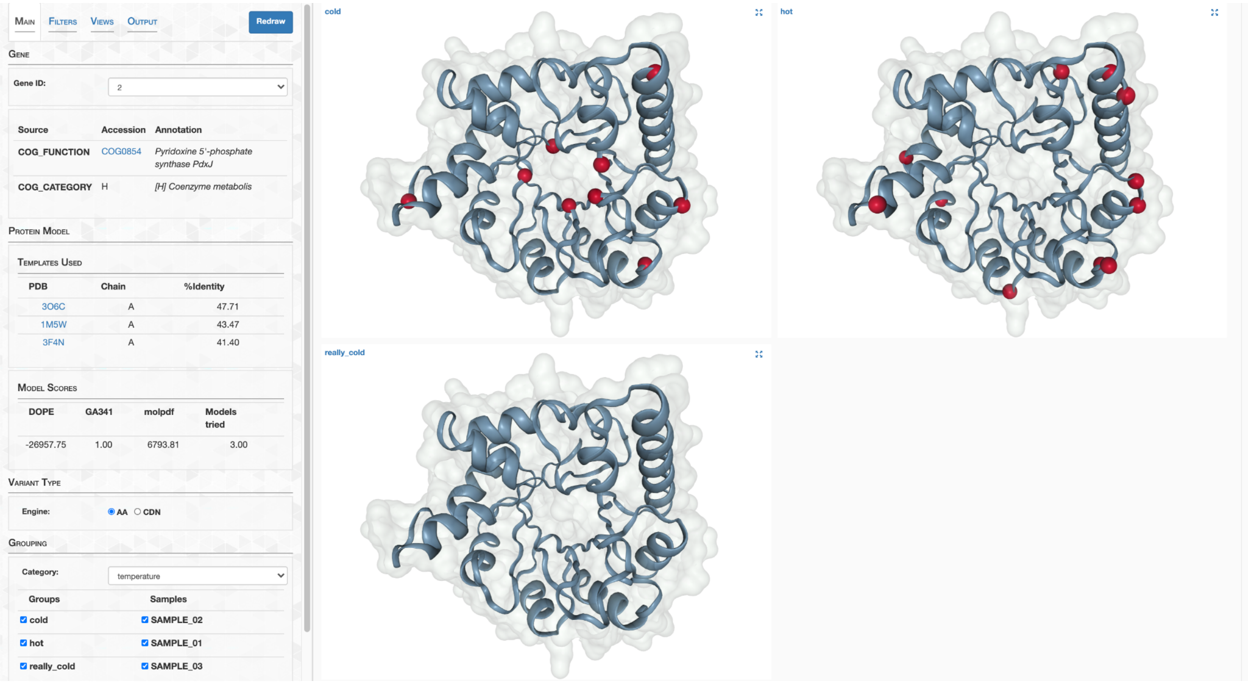


**Figure SI7. Screenshot of the interface with the "Main" tab active.** The user has chosen to visualize Gene ID 2 from the left-hand side panel. Functional annotations from COG indicate this is a Pyridoxine 5’-phosphate synthase, and its structure was modeled using the PDB IDs 3O6C, 1M5W, and 3F4N templates. The resulting structure is visualized on the right-hand side in 3 separate views corresponding to each of the 3 groups of metagenomes specified by the user in the bottom left corner. The spheres overlaid onto the 3 views are the positions of single-amino acid variants found from each group, and can be switched to single-codon variants by switching the Variant Type Engine from “AA” (amino acid) to “CDN” (codon).

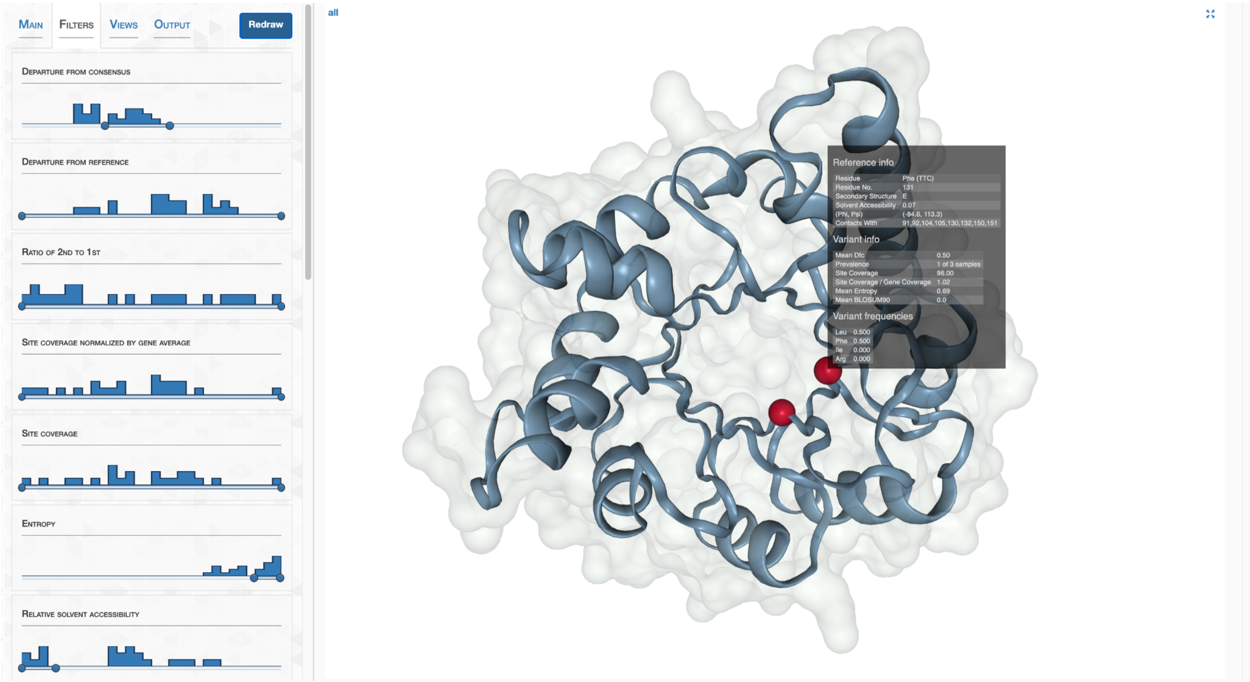


**Figure SI8. Screenshot of the interface with the "Filter" tab active.** Variants can be filtered in the “Filters” tab, which shows a suite of filters, each represented as an interactive slider with endpoints that can be clicked and dragged by the user. Above each slider is a histogram detailing how the variants distribute according to the filter. In this screenshot, the user has included variants with mid-range “departure from consensus” values, high “entropy” values, and low “relative solvent accessibility” variants. The right-hand side reveals that two variants (red spheres) match this filter criteria. Hovering the mouse above one of the variants activates a pop-up menu from which relevant statistics can be learned about.

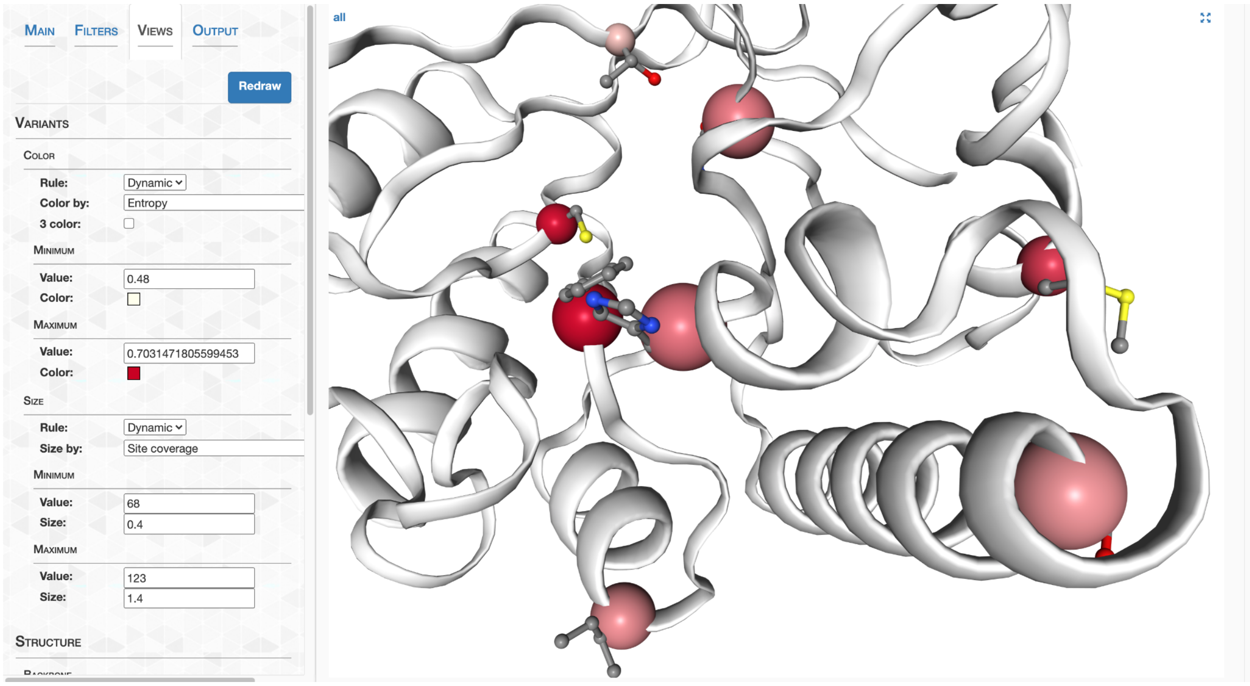


**Figure SI9. Screenshot of the interface with the "Views" tab active.** In the “Views” tab, variants can be colored and sized according to variables. In this screenshot, the user has colored variants according to their entropy values on a linear gradient between white and red, and sized them according to their metagenomic coverage values.

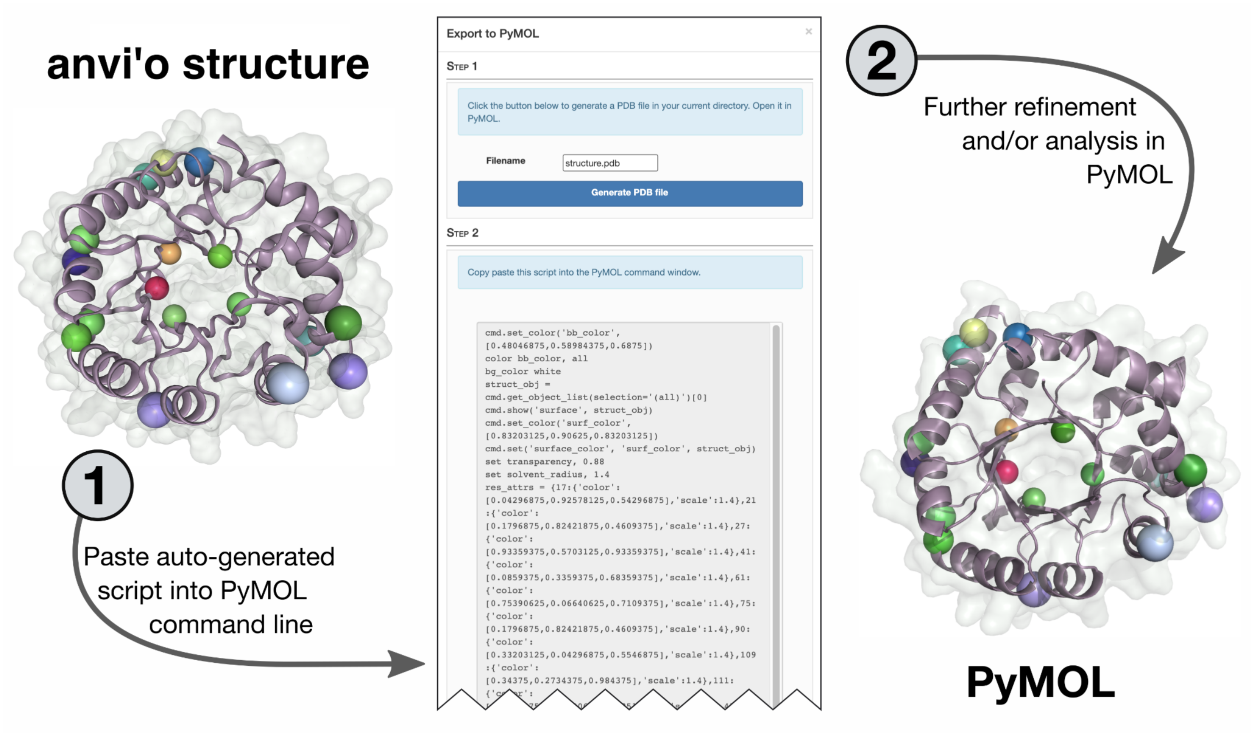


**Figure SI10. The interface can seamlessly migrate user sessions into PyMOL for visual refinement and more sophisticated analysis than is possible with `anvi-display-structure`**. Under the "Output" tab, users can select "Generate in PyMOL" to auto-generate a script (middle) that when pasted into the PyMOL command line, reproduces the current interface view directly in PyMOL.

Edgar, Robert C. 2004. "MUSCLE: A Multiple Sequence Alignment Method with Reduced Time and Space Complexity." *BMC Bioinformatics* 5 (August): 113.

Jumper, John, Richard Evans, Alexander Pritzel, Tim Green, Michael Figurnov, Olaf Ronneberger, Kathryn Tunyasuvunakool, et al. 2021. "Highly Accurate Protein Structure Prediction with AlphaFold." *Nature* 596 (7873): 583–89.

Rose, Alexander S., Anthony R. Bradley, Yana Valasatava, Jose M. Duarte, Andreas Prlić, and Peter W. Rose. 2016. "Web-Based Molecular Graphics for Large Complexes." In *Proceedings of the 21st International Conference on Web3D Technology*, 185–86. Web3D '16. New York, NY, USA: Association for Computing Machinery.

Rose, Alexander S., and Peter W. Hildebrand. 2015. "NGL Viewer: A Web Application for Molecular Visualization." *Nucleic Acids Research* 43 (W1): W576–79.

Worth, Catherine L., Sungsam Gong, and Tom L. Blundell. 2009. "Structural and Functional Constraints in the Evolution of Protein Families." *Nature Reviews. Molecular Cell Biology* 10 (10): 709–20.

Xu, Jinrui, and Yang Zhang. 2010. “How Significant Is a Protein Structure Similarity with TM-Score = 0.5?” *Bioinformatics* 26 (7): 889–95.

Zhang, She, James M. Krieger, Yan Zhang, Cihan Kaya, Burak Kaynak, Karolina Mikulska-Ruminska, Pemra Doruker, Hongchun Li, and Ivet Bahar. 2021. “ProDy 2.0: Increased Scale and Scope after 10 Years of Protein Dynamics Modelling with Python.” *Bioinformatics* , April. https://doi.org/10.1093/bioinformatics/btab187.

Zhang, Yang, and Jeffrey Skolnick. 2004. “Scoring Function for Automated Assessment of Protein Structure Template Quality.” *Proteins* 57 (4): 702–10.
